# Conformational dynamics of a histidine molecular switch in a cation/proton antiporter

**DOI:** 10.1101/2024.12.20.629634

**Authors:** Cristina Pecorilla, Anton Altmeyer, Outi Haapanen, Yongchan Lee, Volker Zickermann, Vivek Sharma

## Abstract

Multisubunit Mrp type sodium proton (Na^+^/H^+^) antiporters are indispensable for the growth of alkali and salt tolerant bacteria and archaea under challenging conditions. Several subunits of the membrane protein complex are closely related to the membrane bound subunits of mitochondrial respiratory complex I, a key enzyme in aerobic energy metabolism. The molecular mechanism of ion translocation by complex I and Mrp antiporters has remained largely unknown and is the subject of intense debate. Here, we combine the power of site-directed mutagenesis and large-scale atomistic molecular dynamics simulations to elucidate the molecular basis of proton-conducting function of the MrpA subunit of the antiporter. We show that point mutations directly affect the transport activity by perturbing the conformational dynamics of a key histidine residue that acts as a molecular switch. Importantly, we find that charge and protonation state variations drive hydrogen bonding rearrangements and hydration changes, which result in coupled sidechain and backbone level conformational changes of the histidine and neighboring residues. We propose the histidine switch as a unique functional element central to proton transport function in Mrp antiporters and respiratory complex I.

**Significance statement:** Energy conversion in biology is catalyzed by several membrane-bound enzyme complexes that drive transmembrane charge translocation. A key question is how these charges are moved against a membrane gradient in an efficient manner, i.e., what kind of gating and fail-safe mechanisms are employed by the proteins to ensure charge transfer directionality. By studying cation/proton Mrp antiporter, which shares homology with the mitochondrial complex I, we describe a conserved histidine residue acting as a molecular switch crucial for gating transfer of protons. Our work emphasizes the functional significance of conserved and conformationally mobile motifs in proteins. The results will benefit the mechanistic understanding of naturally existing proteins and design of artificial enzymes.

## Introduction

Sodium proton (Na^+^/H^+^) antiporters are conserved across all kingdoms of life and are indispensable for intracellular pH homeostasis and salt tolerance^1-4^. In well studied sodium proton antiporters of the monovalent cation proton antiporter family (CPA1 and CPA2), the antiport activity is associated with a single transmembrane protein^5-8^. In contrast, the Mrp (multiple resistance and pH adaptation) type Na^+^/H^+^ antiporters are large multisubunit complexes, in which all subunits appear to be relevant for function^9,10^. The Mrp type antiporters are widely distributed in bacteria and archaea and facilitate sodium and pH homeostasis under challenging conditions. Mrp antiporters have been identified as critical for the pathogenicity of bacteria such as *Staphylococcus aureus* and *Acinetobacter baumanii* that are major threats to human health^11,12^. While Mrp antiporters evidently play an important role in bacterial physiology, they are also the focus of current research because they have an evolutionary relationship to respiratory complex I and other members of the complex I superfamily^13-16^. Recently, three structures of Mrp type antiporters have been determined by cryo electron microscopy (cryo-EM) and different models for proton and sodium translocation pathways have been proposed^17-19^. Our structure of the seven-subunit antiporter from *Bacillus pseudofirmus* (Fig. 1) has the best resolution to date and allowed modeling of more than 300 water molecules^19^. About 70 protein bound waters are arranged along a central hydrophilic axis that is highly similar to the situation found in the membrane arm of complex I. With 21 transmembrane helices (TMHs), MrpA is the largest subunit and has an unusual topology (Fig. 1). It can be divided into spatially separated N-terminal (14 TMHs) and C-terminal (5 TMHs) domains that are connected by a long horizontal helix with two anchoring TMHs. There is a consensus that the N-terminal domain of MrpA is responsible for proton uptake from the periplasm and transfer to the cytoplasm^17-19^. We and others have proposed that sodium binding and transfer occurs remote from this pathway and involves several residues of the C-terminal domain of MrpA^17,19-21^. It follows that in contrast to single subunit antiporters, but in striking similarity to complex I, a long-range coupling mechanism has to be envisaged for Mrp type antiporters.

**Figure 1.**
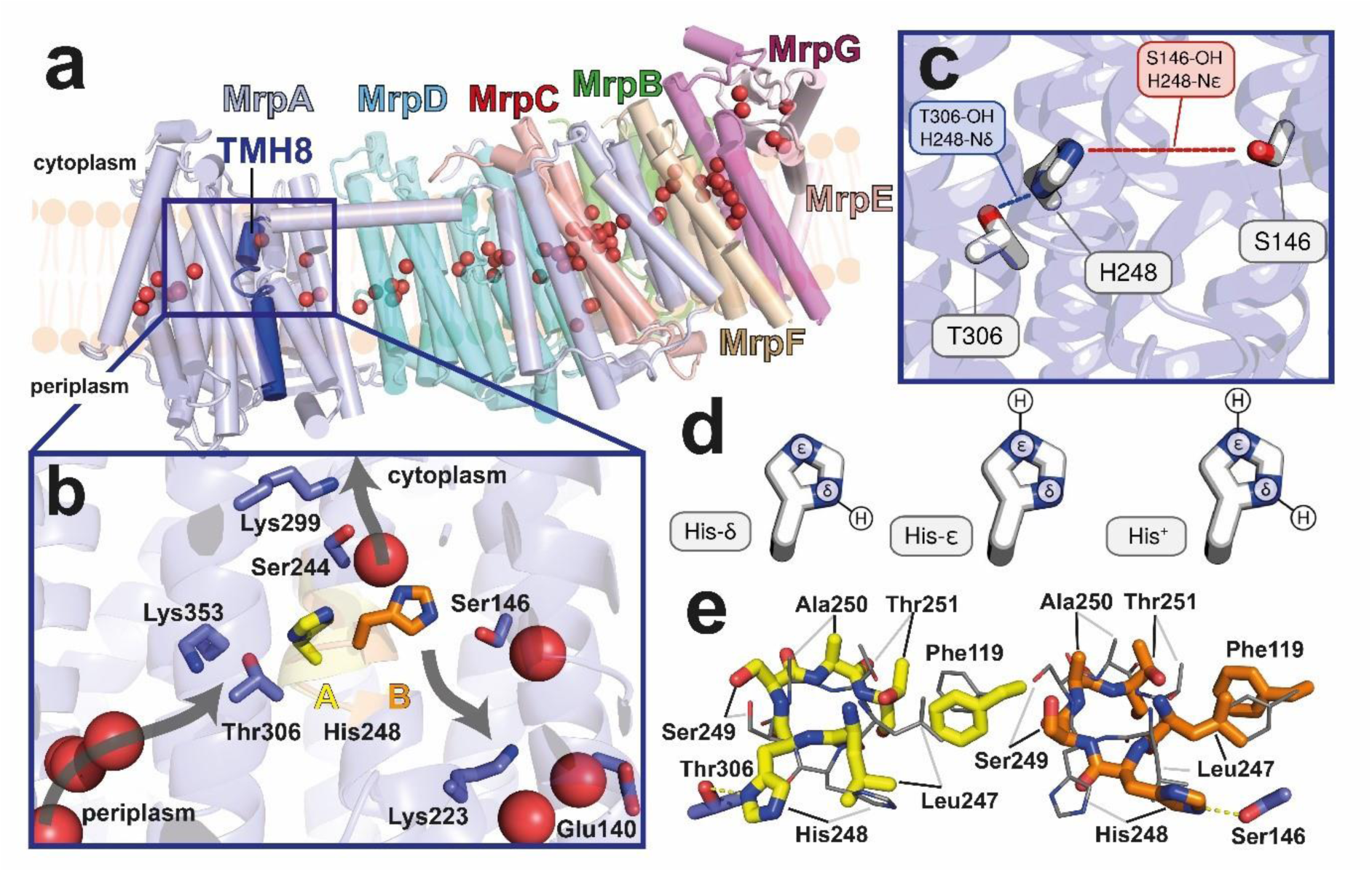
Architecture of Mrp-antiporter and histidine switch. **(a)** Side view of the cryo-EM structure of the seven subunit Mrp type sodium/proton antiporter from *B. pseudofirmus* (PDB 7QRU); protein bound water molecules (red spheres) form a hydrophilic axis in the core of the protein complex, TMH8 (dark blue) in the N-terminal domain of the MrpA subunit is interrupted by a mobile segment. **(b)** H248 sits in the mobile segment of TMH8 and at the junction of three putative proton translocation pathways (arrows show the potential proton transfer directions). In the A conformation it forms a hydrogen bond with T306 and in the B conformation it forms a hydrogen bond with S146. **(c)** Distances between imidazole nitrogens and hydroxyl oxygens of T306 and S146 are marked to visualize the mobility and preferred orientation of H248 in MD simulations (see below Fig. 3). **(d)** Possible protonation states of the histidine side chain. **(e)** View from cytoplasmic side onto the mobile segment (residues 247-251) of TMH8 (A conformation, left, yellow; B conformation, right, orange; overlay with the other conformation is shown in gray); F119 from TMH4, a mitochondrial disease locus in ND5 subunit of complex I is thought to limit the mobility by blocking further movement of the sidechain of L247 in the B conformation.

The fold of the N-terminal MrpA domain is clearly related to the fold of the neighboring MrpD subunit. Consistently, TMH7 and TMH12 are interrupted by short loop segments, a strictly conserved glutamic acid residue is present in TMH5 and strictly conserved lysine residues sit in TMH7, TMH8 and TMH12. An important difference is that a strictly conserved histidine residue (H248; all amino acid numbering corresponds to MrpA subunit of *B. pseudofirmus*, see Fig. 1) is present in a partly α helical region of TMH8 of MrpA but absent in MrpD. While MrpA shares a plethora of conserved features with mitochondrial complex I subunit ND5, including the conserved TMH8 histidine, the MrpD subunit is more closely related to complex I subunits ND2 and ND4.

Atomistic molecular dynamics (MD) simulations of bacterial respiratory complex I identified three conformational states of the conserved TMH8 histidine^22^. Several cryo-EM structures of complex I have resolved the conserved histidine in different conformations^23-26^, highlighting consistency between structural and simulation data. The cryo-EM maps of the *B. pseudofirmus* antiporter provided strong evidence that the seven-residue segment of TMH8 with strictly conserved H248 at its center is mobile and adopts two alternative conformations (Fig. 1)^19^. In alternative conformation A, the histidine points towards strictly conserved T306 and connects to a putative proton uptake pathway from the periplasm. In the B conformation the histidine points towards strictly conserved S146 on a putative pathway to the canonical TMH7 K223/ TMH5 E140 pair (Lys/Glu pair) of MrpA that is part of the central hydrophilic axis. In the B position, H248 also connects via a water molecule to strictly conserved S244 on a putative pathway to the cytoplasm. The critical role of H248 was supported by complete loss of antiport activity in a H248A mutant and MD simulations indicated that the position of the histidine sidechain is closely connected to its protonation state^19^. Based on these findings and in line with the previous simulations of H248 and the corresponding region in complex I^22^ we proposed that the histidine residue enables gated transmembrane proton transfer in the MrpA subunit of the antiporter and the ND5 subunit of complex I^19,27^.

In this work, we performed site-directed mutagenesis of several conserved amino acid residues in the region around H248, including amino acid substitutions that emulate mitochondrial disease mutations in complex I. To obtain molecular insights into the wild type and mutant proteins’ function and dysfunction, respectively, ca. 150 microseconds of atomistic MD simulations are performed on our high-resolution structure of the Mrp antiporter in several different protonation and conformational states. The powerful approach of combining extensive mutagenesis of the MrpA subunit with the large-scale atomistic MD simulations provide a detailed mechanistic model of the proposed histidine switch mechanism in Mrp antiporter and respiratory complex I.

## Results

### Function of selected residues analyzed by site-directed mutagenesis

Based on our cryo-EM structure^19^, we identified 10 residues of interest in close proximity to H248 and introduced a total of 20 single-site mutations in the MrpA gene that modify the polarity and conformational flexibility of the region. All of the selected residues are highly conserved in Mrp antiporters and respiratory complex I^19^ and encompass the mobile segment of TMH8 as well as the three polar residues (T306, S146 and S244) that connect H248 to three putative proton transfer pathways (Fig. 1). In addition, we investigated the role of conformationally mobile F119 in the adjacent TMH3 (Fig. 1), whose counterpart in complex I has been linked to Leigh disease^28^, and of the strictly conserved W232 in the loop region of discontinuous TMH7.

To characterize the functional effects of residue exchanges, we expressed mutated variants of the *B. pseudofirmus* Mrp antiporter in the *Escherichia coli* strain KNabc(DE3), that is deficient in the three canonical Na^+^/H^+^ antiporters of *E. coli* (NhaA, NhaB and ChaA)^29^. The expression level was monitored immunologically and did not show considerable differences between the wild type Mrp antiporter and the majority of mutants (Fig. S1). We found a partial reduction in expression level for mutants H248A and T306S and an about twofold increased level of expression for mutants F119L, W232A and A250C. Baseline KNabc(DE3) cannot support growth beyond 200 mM NaCl, while supplementation of KNabc(DE3) with the wild type Mrp antiporter from *B. pseudofirmus* sustains growth in medium with up to 800 mM NaCl. We further queried the effect of selected residue exchanges on the activity of the antiporter with a fluorescence based dequenching assay in inverted membrane vesicles of KNabc(DE3) (see Methods and Fig. 2d)^30^. All mutants could be clustered in groups following the behavior of either negative control (Fig. 2a), wild type (Fig. 2c) or displaying an intermediate phenotype (Fig. 2b).

**Figure 2.**
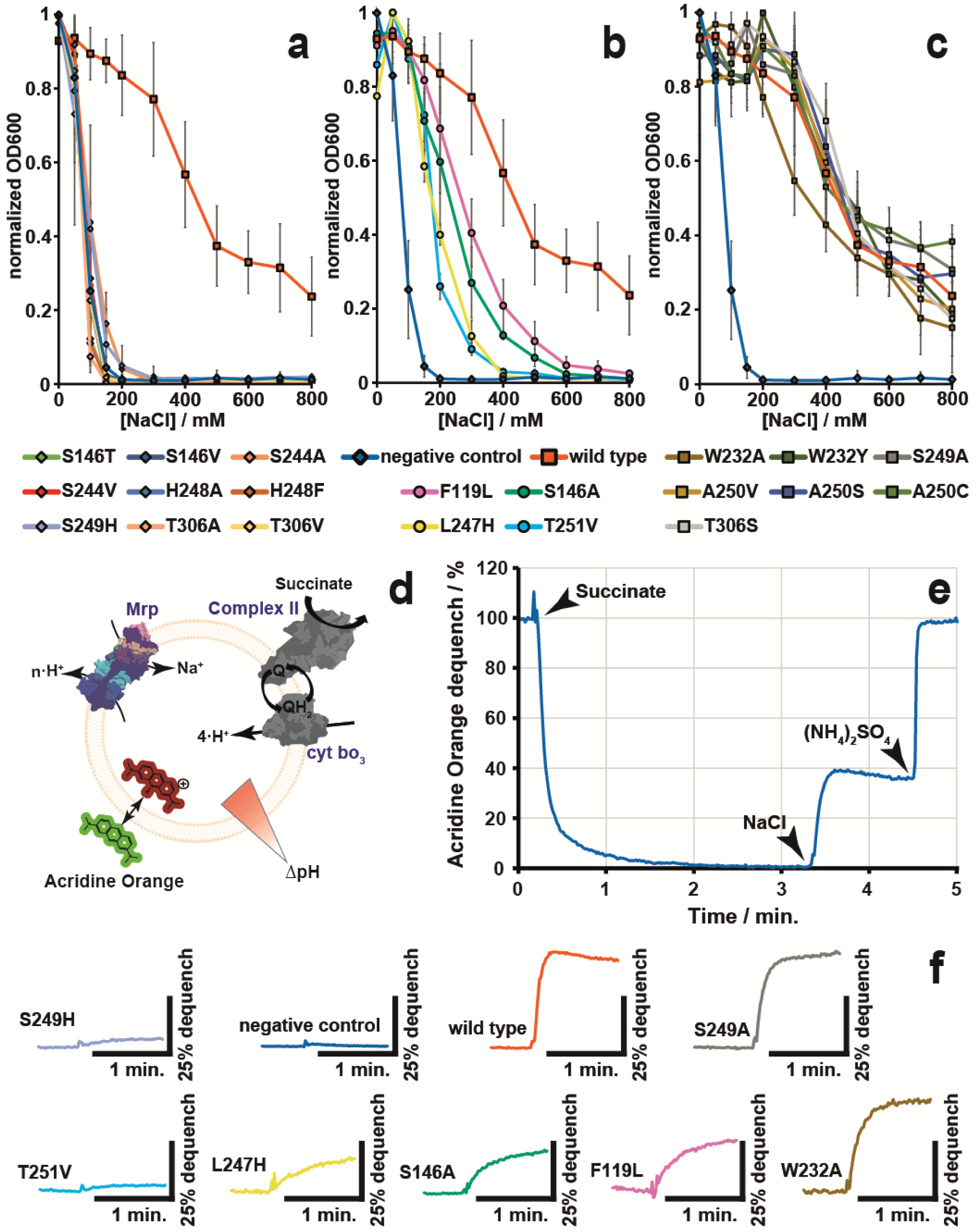
Effects of mutants related to histidine switch mechanism on salt resistance and antiport activity. (**a-c**) Test for ability of mutants to confer salt resistance to the *E. coli* triple Na^+^/ H^+^ antiporter knock-out strain KNabc(DE3). Mutants that confer salt-resistance on the level of the negative control (here the empty plasmid pUB26) (a) are marked with diamond shapes, while mutants that confer salt-resistance on similar levels to wild type protein (c) are marked with square shapes. Intermediate level of salt-resistance (b) is marked with circular shapes. (**d**) Cartoon depicting the setup for antiport measurements in inverted KNabc(DE3) membrane vesicles. Addition of succinate causes part of the respiratory chain consisting of Complex II (based on PDB: 1NEN) and cytochrome *bo*3 (based on PDB: 7CUB) to acidify the inside of the vesicles. The fluorescent probe acridine orange (AO) changes its emission wavelength within acidified vesicles. Addition of sodium allows Mrp antiporter (based on PDB: 7QRU) to extrude protons in exchange for sodium uptake. (**e**) Example trace of an antiport measurements for wild type protein. The injection events are marked and labeled. Addition of 3.75 mM succinate results in quenching of AO, while addition of 10 mM sodium causes the AO to dequench by about 40 %. The addition of 8 mM ammonium sulfate abolishes the proton gradient, resulting in full dequenching of AO. (**f**) Partial traces of antiport measurements for chosen mutants, wild-type, and empty plasmid. Only steady state region before the addition of sodium and the sodium dequenching are shown. Colors correspond to a-c.

To start, we reviewed the role of H248 by mutating it to alanine and bulkier phenylalanine. In both cases and in agreement with our previous study^19^, the antiporter lost its ability to confer salt resistance (Fig. 2a). Next, we sought to create a situation in which alternative positions A and B are simultaneously occupied by histidine residues by mutating S249 or L247 to this residue. We hypothesized that this would restrict the conformational mobility of H248 in the S249H and L247H mutant and the antiporter would lose its activity. In the case of the S249H mutant, indeed the ability to confer salt resistance and dequenching activity was entirely lost (Figs. 2a and 2f), while surprisingly in the case of L247H we observed an intermediate phenotype, that supported growth only up to 300 – 400 mM NaCl (Fig. 2b) and still showed partial dequenching activity (Fig. 2f). Mutation of S249 to alanine on the other hand resulted in wild type-like behavior (Figs. 2c and 2f).

We next analyzed the three polar residues that participate in three putative proton transfer pathways shown in Fig. 1b. The residue T306 that connects to the periplasmic path was mutated to alanine, serine, and valine. Both exchanges that remove the hydroxyl group, T306A and T306V, cause loss of Mrp antiporter mediated salt resistance (Fig. 2a). In contrast, the mutation that preserves the polar moiety, T306S, displayed wild type-like behavior (Fig. 2c). This indicates that the polar moiety is essential in this location. A more intriguing picture emerged upon mutation of the serine residue (S146) that participates in the pathway connecting to the Lys/Glu pair and MrpD (Fig. 1), and is also conserved in respiratory complex I. Here we mutated the residue S146 to alanine, threonine, and valine. While the mutant S146V removed the ability to confer salt resistance, similar to its counterpart T306V, so did the mutant S146T, despite conserving the polar moiety (Fig. 2a). The mutant S146A on the other hand displayed another intermediate phenotype, this time supporting the growth up to 500 – 600 mM NaCl (Fig. 2b), and showed residual dequenching activity (Fig 2f). This is surprising given that S146 forms a hydrogen bond with H248 in the B conformation. For the pathway connecting H248 to the cytosol, we mutated the residue S244 to alanine and valine. In both S244A and S244V, the loss of the polar moiety removed the ability to confer salt resistance (Fig. 2a), indicating that the polar residue as well as the water molecule bound by this residue are important for antiporter function.

We followed up by querying the residue T251, which is strictly conserved in complex I and highly conserved in Mrp antiporters. In our cryo-EM structure, the hydroxyl-group of its sidechain is involved in hydrogen bonding interactions with the backbone oxygen atom of L247 (Fig. 1e, see below), indicating that it might be involved in stabilizing the non-α helical character of the mobile region (see Fig. 1 and Introduction). We therefore removed the hydroxyl-group by mutating it to a non-polar valine. The mutant T251V displayed a similar growth phenotype to L247H, supporting growth up to 300 – 400 mM NaCl but showed almost no dequenching activity indicating an important functional role for this residue (Figs. 2b and 2f).

A250 is part of the mobile region of TMH8 (Fig. 1). The corresponding residue in human complex I is a serine and its mutation to cysteine has been described as a disease mutant^31^. We reasoned that possibly a small residue is important at this position but mutation of the alanine to valine, serine or cysteine in MrpA did not have any clear negative consequences (Fig. 2c). Although the higher expression level of mutant A250C could in principle conceal a hypothetical decrease of Mrp antiporter function, the fact that position 250 does not respond to two other exchanges suggests that the pathogenic effect is specific to human complex I. Besides S250C (human complex I numbering), there is another disease mutant in human complex I in the corresponding region of the ND5 subunit. The mutation F124L (human complex I numbering) has been confirmed to be involved in Leigh disease^28,31^. The corresponding residue in MrpA is F119, which sits in TMH3 and undergoes moderate conformational changes in step with L247 (Fig. 1e). The leucine mutant F119L displayed an intermediate phenotype supporting growth up to 600 – 700 mM NaCl (Fig. 2b) and decreased dequenching. This indicated that the mutant has a considerable effect on the mechanism that is conserved up to mammalian complex I. We note that F119L mutant may have an even stronger negative effect on antiport function, which could be masked by the higher expression level of this mutant.

Finally, we tested the strictly conserved aromatic residue W232, which sits in the loop segment interrupting TMH7 and points towards the mobile TMH8 element (Fig. S26a). We reasoned that it might play a role in controlling the movement of the nearby residues, including H248. We therefore mutated the residue to alanine and tyrosine. Both the removal of the bulky sidechain with W232A and the introduction of a large polar moiety with W232Y resulted in wild type-like behavior (Figs. 2c and 2f) excluding a critical role of this residue in antiport activity, albeit a minor reduction in Mrp antiporter function may not be recognizable due to its higher expression levels.

### All-atom MD simulations of wild type Mrp antiporter

To obtain a molecular understanding of the site-directed mutagenesis data, we resorted to extensive atomistic MD simulations of the Mrp antiporter in full membrane-solvent environment in multiple charge and conformational states (see computational methods). First, we assessed the main characteristics of the wild type Mrp antiporter simulations, with a focus on H248 and the region around it. We modeled both the neutral protonation states of H248, namely H248ε and H248δ, as shown in Fig. 1d. The protonation states of titratable residues surrounding H248 were fixed based on p*K*_a_ calculations (see computational methods and Table S3). Moreover, because H248 and some residues in its vicinity were structurally observed in the two alternative conformations A and B (see Fig. 1), we initiated MD simulations of H248δ and H248ε protonation states from alternative locations A and B, respectively (see computational methods and Table S4). The first and most straightforward variables to distinguish between A and B positions of H248 are the distances of its imidazole sidechain moiety from T306 and S146, highlighted in Fig. 1c. We find that in the wild type simulations, both A and B conformations can be explored by the H248ε tautomeric state (Fig. 3a). Conversely, H248δ populates only the A conformation with somewhat lower occupancy (Fig. S2). We note that in the A position, T306 makes a hydrogen bond to H248, and acts as a hydrogen bond donor in the H248ε state. In the B conformational state of histidine, S146 makes a stable hydrogen bond to H248, and in this case histidine acts as a hydrogen bond donor to serine. These hydrogen bonding arrangements of H248 with the polar residues are likely to be important for the proton transfer pathways (Fig. 1, see more below).

**Figure 3.**
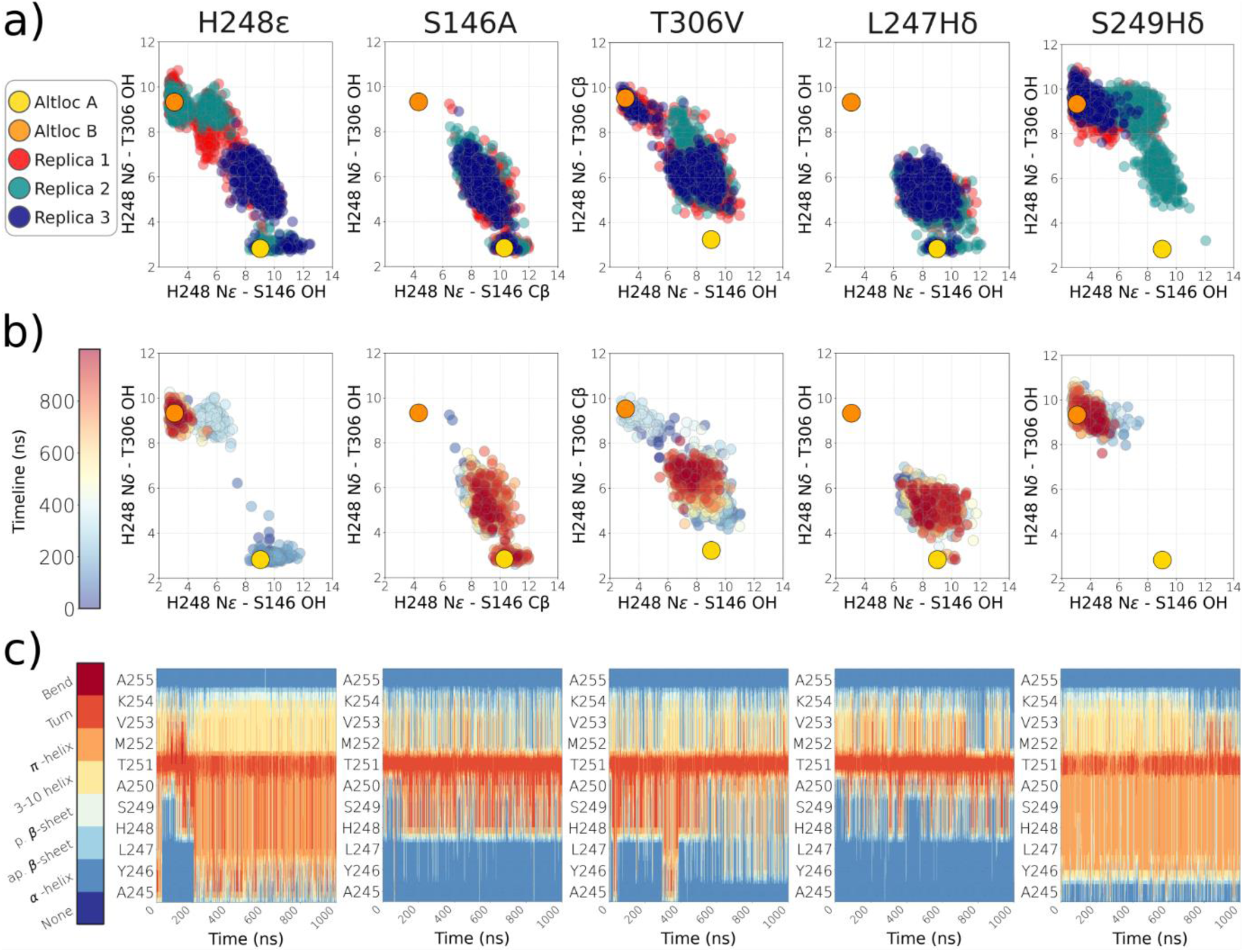
Conformational dynamics of the sidechain of H248 and the backbone carrying it in MD simulations of wild-type and mutant states. **(a)** Scatter plots displaying the distances of S146 and T306 side chains from H248ε side chain, where distances are pictured in Fig. 1c (for H248δ, see Supplementary Figures). X axis measures the distance between ε nitrogen atom of H248 and S146 oxygen (Cβ for S146A). Y axis measures the distance between δ nitrogen atom of H248 and T306 oxygen (Cβ for T306V). Data is shown from all three simulation replicas in three different colors. Note, all H248ε state simulations were initiated from alternative conformation B, whereas H248δ simulations from alternative conformation A (see SI). Yellow and orange spheres mark the position of H248 sidechain (from S146 and T306) in conformation A and B, respectively, as observed in the cryo-EM structure. **(b)** Time resolved behavior of H248 sidechain from selected simulation replica from time 0 (blue) to 1 µs (red). **(c)** Time evolution of the secondary structure profile of H248 carrying segment, corresponding to simulation replica shown in (b).

Interestingly, besides the two clusters corresponding to A and B states observed in MD simulations, the histidine sidechain also explores intermediate stable states that occupy the region in between (Fig. 3). The intermediate conformation has been observed in the MD simulations of respiratory complex I^22^ and an AlphaFold^32^ model of MrpA also captured this intermediate state (Fig. S26b), which may be of functional importance, as discussed below.

Next, we expanded the view from the rearrangements of the H248 sidechain to the conformational changes in the entire helical segment around this residue. Because part of the helix spanning from A245 to A255 shows a non-canonical helical arrangement (Figs. 1 and S27), we performed the secondary structure analysis on simulation trajectories. As shown in Fig. 3, the TMH8 histidine when modeled in H248ε state can populate both the states A and B, with a concomitant change in the secondary structure profile of the segment. The A conformation shows a higher α-helical character, while in the B conformation helicity is largely lost (Fig. 3c and S2, see also Fig. S27). Notably, we do not observe such a change for the other tautomer because H248δ does not populate the B state in simulations (Fig. S2). With these results at hand on wild type system, we next investigated the effects of point mutations (Fig. 2) on H248 conformation, secondary structure of the segment carrying it, as well as hydration in the region, and compared the results with the wild type simulations.

### Molecular insights on antiporter dysfunction by atomistic MD simulations

The polar residues T306 and S146 participate in two of three putative proton transfer pathways in MrpA and form hydrogen bonds with H248 in its alternative locations A and B, respectively (Fig. 1). We therefore first modeled the mutations T306V and S146A/T to observe how H248 dynamics is affected upon removal of a hydrogen bonding partner. As shown in Figs. 3b and 3c for H248ε (Fig. S3 for H248δ), mutation T306V eliminates the population of the A conformation. For both tautomeric states of histidine, the distance between H248 and valine in T306V mutant is found to increase concomitant with the lower occupancy of state A (Figs. 3, 4b). The loss of the A conformation is likely to alter the ability of the Mrp antiporter to transfer protons from the periplasmic side (Fig. 1), and is in agreement with the impaired antiport function seen in experiments (Fig. 2). Similarly, in the case of the S146A mutation, we observe a clear loss of B conformation (drop from 40% to zero, see Figs. 3, 4b, S4), commensurate with the drop in antiporter sodium tolerance (Fig. 2). Interestingly, the more conservative S146T mutation also shows an absence of the B conformation (Fig. S5), indicating that the presence of a methyl group reduces the capability of H248 to participate effectively in hydrogen bonding with threonine in this position. This loss of B population is also clearly observed in secondary structure analysis (Figs. 3, S4 and S5). A lower but still observable sodium tolerance and dequenching activity observed in the S146A mutant (Fig. 2) is likely to be related with the subtle difference between threonine and alanine, where the short sidechain of the latter may create more space for transient water molecules compensating for the loss of serine. Overall, the above data from mutant MD simulations (performed in wild type charge state, cf. below) suggests that by removing the hydrogen bonding partners of H248, the capability of the histidine to adopt the two structurally characterized conformations is impaired.

**Figure 4.**
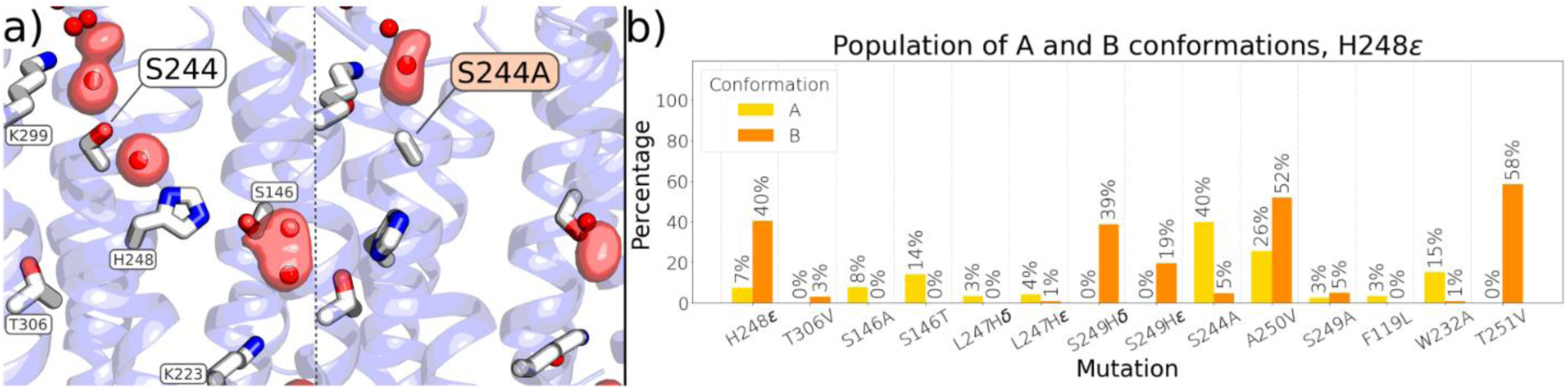
Mutations drive hydration and conformational changes. **(a)** Difference in hydration in the region around H248 in wild type (left) and S244A (right) mutant. Water occupancy map shown in the red glass isosurface is calculated over a total of 3 µs trajectory data using oxygen atoms of water molecules and is shown at an isovalue of 0.2 (20% occupancy). **(b)** A and B conformational occupancies in wild type and mutant conditions. A conformation is adopted when the distance between H248-Nδ and T306-Cβ is below 3.75 Å, whereas B conformation is adopted when the distance between H248-Nε and S146-Cβ is below 4.75 Å. Corresponding analysis for H248δ available in Fig. S30.

To further probe the conformational dynamics of the helical region carrying H248, we proceeded with the mutations L247H and S249H. Both these mutations cause activity loss in experiments, albeit to a varying extent (Fig. 2). In our simulations, we find that with these mutations H248 can no longer transit from state A to B and *vice versa* (Figs. 3 and S6-9). In mutant L247H, the histidine at position 247 replaces H248 in the B conformation, forcing the native histidine into a conformation relatively closer to A with a similar occupancy as wild type (3% vs 7%, Fig. 4b). On the other hand, H248 is locked close to its position in the B conformation in the S249H mutation, with a similar occupancy as in wild type (Figs. S8, S9, see also 4b). In this state, the mutated residue does not move from its original position and prevents the rotation of the neighboring native histidine. This sidechain locking is also reflected on the secondary structure of the helical segment; L247H shows higher α-helical character, whereas S249H shows a strong loss of helicity for both tautomeric states of H248 (Figs. 3 and S6-9). These results, both at the sidechain and backbone levels, are consistent with the data from our wild type simulations performed in same charge state, and highlight that the flexibility of the helical segment plays a crucial role in antiporter function. Even though both the exchanges limit the mobility of H248, scrutinization of simulation trajectories reveal clear differences between the two mutants, which provides an explanation to the differences observed in sodium tolerance and dequenching experiments (Fig. 2). Remarkably, we find that a bridging water-based connection emerges in the L247H mutant, forming a pathway from T306 towards S146 involving H248, L247H and intervening water molecules (Fig. S32A). This pathway can in part compensate for the loss of conformational flexibility of H248, and explain the non-zero salt tolerance and dequenching observed in L247H mutant (Fig. 2). Importantly, no such bridging is observed in S249H mutant. Second, H248 predominantly occupies the B conformation (like wild type but with zero A state occupancy, Fig. 4b), whereas the histidine at position 249 (S249H) does not substitute for the native histidine A conformation (see Fig. S32), which is central for proton transfer from the periplasmic side (Fig. 1). Furthermore, we notice a hydrated region emerges in the vicinity of S249H mutation and in between TMH3 and TMH11 (Fig. S32), which is likely to cause uncoupling due to the loss of periplasmic protons to the cytoplasmic side. On the other hand, no such hydrated region forms in L247H simulations (Fig. S32), and proton accepting position of H248 is clearly stabilized. These differences speak for the weak but non-zero sodium tolerance and dequenching observed for L247H mutant (and also S146A) and strengthens the notion of importance of A alternative location in proton transfer. In contrast to the severely perturbed dynamics of H248 and the backbone carrying it in several different mutants, the S244A mutation, which results in loss of Mrp antiporter function (Fig. 2), primarily affects the hydration level in the region around H248. As shown in Fig. 4a, the connectivity between S244 and H248 is established by a bridging water molecule, which remains stable during MD simulations, consistent with the cryo-EM data that shows strong density representing a stable water molecule. However, when S244 is mutated to a nonpolar residue, such as alanine, the water molecule is destabilized (Fig. 4a). The lack of a water bridge not only leads to the loss of the hydrogen bond network around H248 but also causes a destabilization of its B conformation (drop from 40% to 5%, and enhancement of A conformations from 7% to 40%, see Fig. 4) as well as a change in secondary structure profile (Fig. S10). Overall, these results suggest that S244 has an important role in stabilizing the hydrogen bonding network around H248, which in turn stabilizes its B conformation. Importantly, in contrast to non-polar exchanges of T306V and S146A that affect putative proton transfers by perturbing A and B conformations respectively, the proton transfer towards the cytoplasmic side is impaired in S244A. These data indicate that polar residues in different proton transfer paths affect antiporter function with different mechanism, but with a common denominator of altered histidine dynamics.

To complement the experiments, we also analyzed H248 mutants that completely eliminated sodium tolerance (Fig. 2). We find that mutations H248A and H248F show a lack of hydration in the region, which is likely necessary for protonic movements towards the periplasmic and cytoplasmic sides of MrpA. In agreement with previous studies^19^, we observe that the water-based connection from S244 to H248 is lacking in these mutants, primarily due to the increased non-polarity of the region (e.g. Fig. 4a). Moreover, the absence of H248 not only impairs the hydration between conserved K353 and ion-pair K223/E140 but also no unique water path is established connecting the two sides (Figs. S26c and S26d). thereby strengthening the notion that switching by H248 is a key mechanism to efficiently bridge three different putative proton transfer pathways. Finally, we tested the effect of T251V and F119L mutations (Fig. 2), where the latter is a mitochondrial disease mutation found in the ND5 subunit of complex I^28,31^. We observed that the T251 sidechain can stably interact with the backbone of L247 in wild type simulations, and the mutation of this residue to a non-polar residue affects the overall stability of the H248 helical section, favoring the B conformation (from 40% to 58%, Fig. 4b) and also affecting the secondary structure profile (Fig. S12). We think that a complete loss of dequenching in T251A is in part explained by a complete loss of A conformation. In contrast to T251V outcome, the B conformation is entirely lost upon F119L mutation, with only a small population of A conformation remaining (Figs. 4b, S13). A similar outcome of loss of B conformation in F119L, S146A and L247H with partial retaining of intermediate/A conformations suggest a common mechanistic mode of dysfunction with which these mutants cause partial loss of sodium tolerance and dequenching (see discussion).

All modeled mutations discussed so far have been found to decrease or completely inactivate antiport capacity of Mrp type antiporter. However, biochemical data shows that A250V, S249A, and W232A display wild type like behavior (Figs. 1, 2 and S26a). In our MD simulations, we find that A250V mutation data show high populations of both A and B states for H248ε (26% and 52%, respectively) in line with the A/B populations observed in the wild type protein, and overall a similar secondary structure response (see Fig. S14). The W232A and S249A mutations display both A and B conformations, albeit with reduced populations (Fig. 4b, see also Figs. S15 and S16).

### MD simulations in different protonation states

We conducted additional MD simulations (Table S4) to investigate how changes in the charge states of nearby titratable residues affect the dynamics of H248 in wild type and mutant conditions. Specifically, we varied the protonation states of three conserved lysine residues – K353, K229, and K223 that encircle the mobile segment containing H248 (Fig. 1, Table S4). Previous studies have shown that mutation of any of the three lysine residues result in a loss of Mrp antiporter activity^18,33^. Furthermore, all three lysine residues are conserved in respiratory complex I of the mitochondrial electron transport chain, thereby highlighting their important functional role across complexes and species.

In the wild type states when all three lysines are modeled charge neutral (Tables S3 and S4), H248ε populates both A and B conformations (7% and 40%, respectively, see Figs. 3 and 4), whereas only the A conformation is occupied in the H248δ state (Fig. S2). In its charge neutral form, K299 assumes a buried conformation and accepts a hydrogen bond from the neighboring S244 (Fig. 5a). However, when its protonated state is simulated (with K353 and K223 modeled neutral), a complete restructuring of hydrogen bonding network takes place, which stabilizes the B conformation of H248 (occupancy changes from 40% to 83%, see Figs. 4b, 5b and 5e) and associated non-helical nature of the segment (Fig. S17). Basically, protonated K299 stabilizes the entire pathway in which H248 adopts the B conformation with a stable hydrogen bond to S146 and with a water mediated hydrogen bond acceptance from S244 (Fig. 5b).

**Figure 5.**
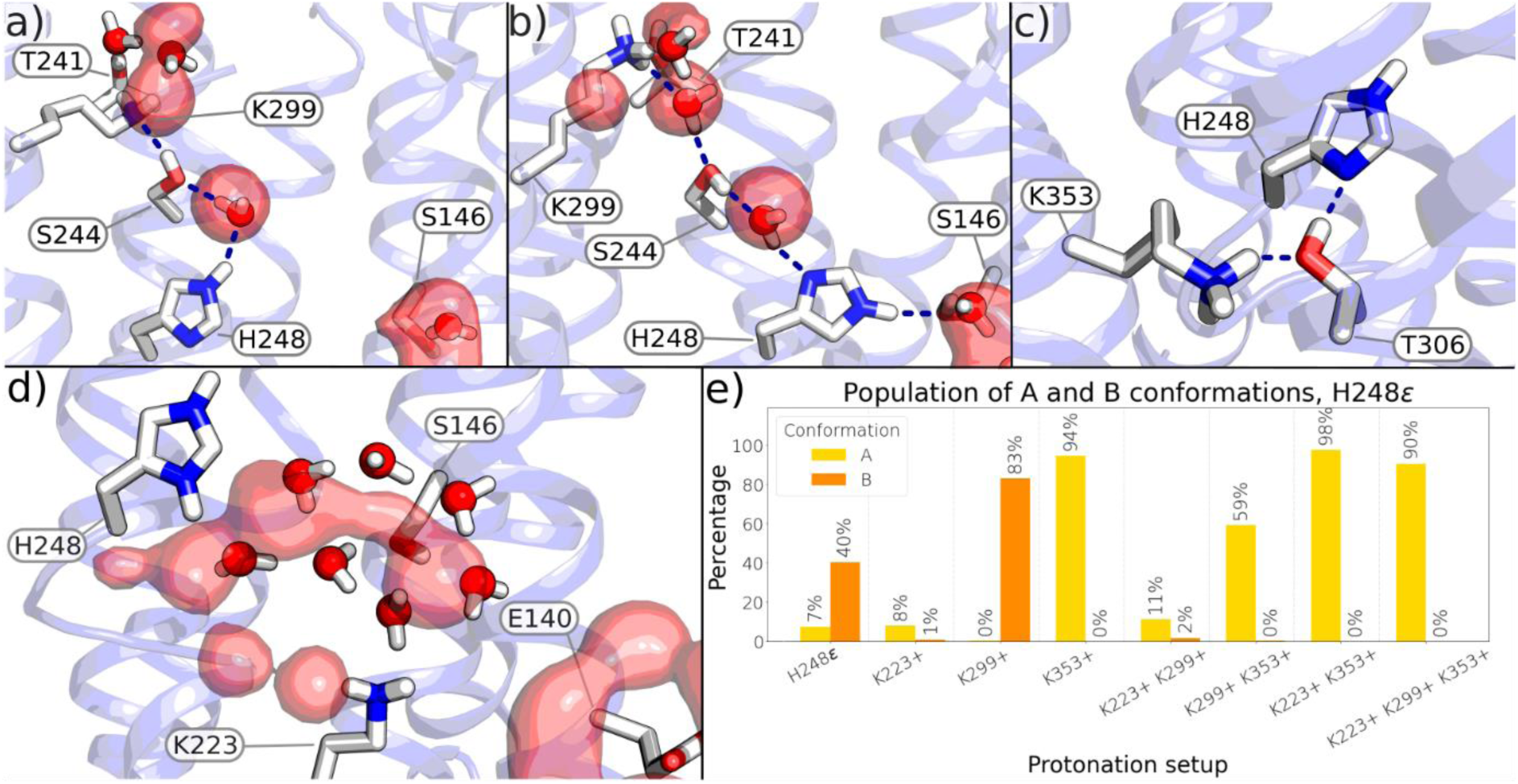
Protonation state changes coupled to hydrogen bonding rearrangements. **(a)** Charge neutral K353 and K299 result in H248ε to adopt different conformations diverse from A and B. Hydrogen bonds are shown with blue dashed lines and water occupancy (red glass isosurface) in panels is shown at an isovalue of 0.2. **(b)** Hydrogen bonding rearrangement takes place upon protonation of K299 resulting in stabilization of the B conformation of histidine. **(c)** Stable hydrogen bonding network between protonated K353 and H248 via T306 stabilizes the A conformation of histidine. **(d)** Positively charged H248 enhances hydration. A new water-based connection is formed between H248 and Lys-Glu pair (neutral K223 and anionic E140). **(e)** A and B conformational occupancies calculated in different charge states of K223, K299, and K353. In H248ε state, all three lysine residues are charge neutral. A conformation is adopted when the distance between H248-Nδ and T306-Cβ is below 3.75 Å, whereas B conformation is adopted when the distance between H248-Nε and S146-Cβ is below 4.75 Å. Corresponding analysis for H248δ available in Fig. S31.

Another striking transition in the sidechain of H248 occurs upon protonation of K353 when it is stabilized predominantly in the A conformation (occupancy up from 7% to 94%, see Figs. 4b and 5e). Scrutinizing the hydrogen bonding pattern between K353, T306 and H248 shows that when the lysine is protonated, it acts as a hydrogen bond donor to threonine, which in turn donates a hydrogen bond to the neutral H248 (Fig. 5c), a scenario that caters for stabilization of the A state as well as enhanced helicity of the TMH segment (Fig. S18). When both K353 and K299 are modeled protonated, the A conformation overrides the B conformation (Fig. 5e, S19), most likely due to the hydrogen bonding described above. All in all, the above data suggest that the A conformation is strongly favored by K353 protonation, while K299 protonation contributes to the stabilization of the B state.

With the knowledge that K299 and K353 are responsible for stabilizing the B and A states, respectively, we tested how mutants S146A and T306V that remove hydrogen bonding partners of H248 alters A/B state occupancies. In agreement with our wild type simulations with K299 and K353 modeled neutral (Figs. 4b and 5e), we observed a drastic loss in B (83% to zero) and A (94% to zero) conformational states of H248 (Fig. S33), thereby strengthening our hypothesis that mutation of hydrogen bonding partners impairs conformational flexibility of histidine, and also affects proton transferring capabilities of the antiporter.

We find that protonation state changes of K223 also affect the H248 conformation, in that it destabilizes the B conformation even if K299 is protonated (Fig. 5e, S20-22). On the other hand, a change in protonation state of K223 has minimal influence on histidine sidechain conformation, as long as K353 is protonated, which stabilizes the A state predominantly (see above).

Besides the variation in protonation state of lysine residues, the histidine was also modeled in its protonated form. In the MD simulations of imidazolium state (His^+^, Table S4) of H248, neither A nor B conformation is observed (Figs. S23, S24 and S26). Instead, the histidine adopts a mid-way position, independently of the protonation state of other residues, and enhances hydration in the region (Fig. 5d). It is the doubly-protonated charge state of histidine that allows a direct water connection of H248 from its intermediate position towards K223 (Figs. 5d and S28b). Notably, the intermediate conformation of H248 resembles the conformation seen in AlphaFold^32^ models (Fig. S26). All in all, our simulation results point out that by perturbing the charged state of H248 as well as of the conserved lysine residues in its vicinity, hydrogen bonding rearrangements take place and conformational dynamics of the central histidine can be altered. This has important mechanistic implications, as discussed below.

## Discussion

One of the major fundamental questions in bioenergetics is how respiratory and photosynthetic enzymes catalyze efficient charge translocation against the membrane potential^34,35^. Respiratory complex I, the first enzyme in the mitochondrial electron transport chain, and bacterial Mrp antiporter, both translocate ions across the membrane by employing long-range coupling^19,36^. To achieve coupling at longer distances, specific structural elements are needed that can enhance the rate of forward reaction and minimize leaks^37-39^. Here, we describe a molecular switch in the form of a conserved histidine residue in the terminal antiporter-like subunits of complex I and Mrp antiporters that may play a key role in gating proton transfer and reinforcing long range coupling. We suggest impairment of the histidine switch mechanism in respiratory complex I is in part responsible for the mitochondrial disease mutations that occur in its ND5 subunit.

The Mrp antiporter couples the transfer of Na^+^ ions from the cytoplasm to the periplasm with proton transfer in the opposite direction. The stoichiometry of electrogenic Na^+^ to H^+^ transport function is unknown, but several proposals have been made^17-19^. In the mode, when protons are exergonically driven from the periplasmic to the cytoplasmic side to catalyze endergonic Na^+^ translocation to the outside, we have earlier discussed a molecular mechanism with the stoichiometry of 2H^+^ consumed against one Na^+^ pumped out^19^. In one of the proton translocation modules, the MrpA subunit, which is homologous to the ND5 subunit of respiratory complex I, a highly conserved histidine residue (H248) has been suggested to undergo conformational dynamics^19,22^. By performing large-scale atomistic MD simulations of bacterial respiratory complex I, multiple conformational states of conserved histidine were identified^22^. Based on high resolution structure and simulations of Mrp antiporter from *Bacillus pseudofirmus*, a mechanism of proton-coupled sodium transport has been proposed^19,27^. In this study extensive site-directed mutagenesis of residues surrounding H248 was performed to probe the effect of mutations on antiport activity. To investigate the molecular aspects of antiporter function and dysfunction, around 150 microseconds of atomistic MD simulations in wild type and mutant states were performed, and backbone and sidechain dynamics of H248 were analyzed. We observed that the removal of H248 or its hydrogen bonding partners as well as the introduction of polar residues in positions vicinal to H248, that stabilizes new hydrogen bonding partners, cause a severe reduction in antiport activity, as observed in growth tolerance analysis and dequenching assays. In addition, the exchange of residues putatively involved with the range of movement for H248 (such as T251V or F119L) cause significant but less severe reduction in antiport activity. In agreement to these findings, we observed that histidine dynamics as part of the histidine switch mechanism can be severely reduced in these mutants.

We emphasize that the conformational dynamics of histidine is affected both by its own protonation state, as well as by the protonation states of several conserved residues in its vicinity, that have previously been shown to severely impact antiport activity^18,33^. Several charge states of the MrpA subunit were studied by varying the protonation states of amino acids together with exhaustive simulation sampling. These data reveal how the protonation state of residues perturb the conformational dynamics of histidine and hydrogen bonding patterns between residues as well as water molecules, and such rearrangements provide directionality to protons to coupling sites as well as to the cytoplasm.

The two alternative conformations of H248 (A and B) are found to be strongly stabilized by the protonation of K353 and K299, respectively. The stabilization is achieved with the reorganization of hydrogen bonding networks. By replacing lysine residues with arginine, which has a higher proton affinity, it may be possible to enhance the stabilization of the hydrogen bonding networks and obtain high enough occupancy of A or B conformations of H242 to be observed in cryo-EM experiments (also on respiratory complex I).

An intermediate state of H248 (other than the A and B alternative conformations) has been suggested to exist based on earlier MD simulations of bacterial respiratory complex I^22^. The occupancy of this state has been observed to be high in T306V and S146A mutant states (also in other mutants) and when the imidazolium state of histidine is simulated, which stabilizes a water-based pathway towards K223 of KE pair. Based on these data, we propose cryo-EM experiments to be performed at low pH and/or in mutant conditions on complex I and Mrp antiporter, which would not only allow trapping H248 in its unique intermediate position but may also provide novel insights on putative proton transfer routes.

Based on the biochemical experiments and molecular simulation data described above, we are now able to propose a detailed model for the histidine switch mechanism (Fig. 6) that is responsible for catalyzing proton translocation in both Mrp antiporter and respiratory complex I. In the ground state of the Mrp antiporter, neutral H248, with its ε nitrogen protonated, hydrogen bonds with S146 and predominantly occupies the B conformation (state 1, Fig. 6). At this stage, K299 is protonated and the conserved K223/E140 pair (Lys/Glu pair) is proton deficient (net charge -1). We note that our MD simulation data suggests that when K299 is protonated and the Lys/Glu pair anionic, the B conformation is preferentially stabilized by means of hydrogen bonding arrangements. The stabilization of the B conformation of H248 in the ground state precludes any protonic connectivity towards the cytoplasmic side as well as to the Lys/Glu pair, which may be important in preventing short circuiting. Indeed, the mutants S146A and L247H, which eliminate the B conformation of H248 entirely (occupancy drop from 40-90% to zero) display loss of sodium tolerance and partial dequenching.

**Figure 6.**
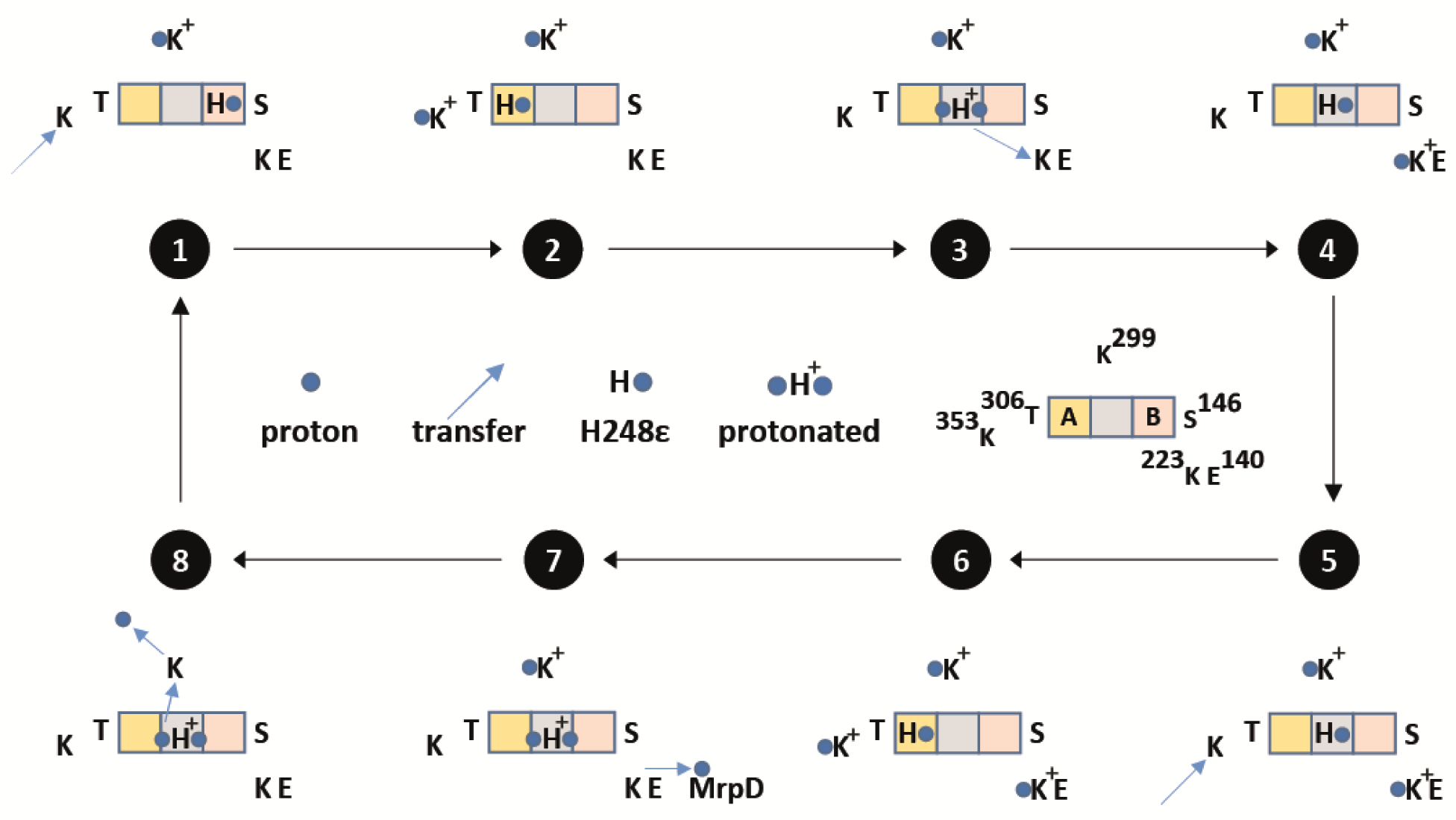
Proposed histidine switch mechanism of proton transport in MrpA subunit. See text for details and Fig. S29 for a proposed mechanism on complex I.

Upon arrival of the first proton to K353 from the periplasm, a remarkable transition of H248 occurs, when it switches its position from B to A conformation (state 2), as consistently observed across several MD simulations. In the A conformation, H248 forms a stable hydrogen bond with T306, which acts as an intermediate proton transfer element from protonated K353 to neutral H248. We note that only in the A conformation H248 accepts a proton arriving from the periplasmic side, thereby, highlighting the key role of this alternative location. The notion is also supported by MD simulation of the inactive T306V mutant in two different charge states that shows a complete loss of A conformation. Upon protonation of histidine, it dissociates the direct hydrogen bonding contact with T306 and assumes an intermediate position (state 3, Fig. 6) – the third location in between the A and B conformations, observed in MD simulations and AlphaFold model. At this stage, protonated H248 has no access to the cytoplasm, but to the conserved Lys of the Lys/Glu pair via a pathway involving water molecules and polar residues, as seen in MD simulations. It is important to note that at this stage of the mechanism, the protonated state of K299 effectively prevents the formation of cytoplasmic connectivity from the protonated histidine, thereby, kinetically preventing the otherwise thermodynamically favorable loss of the proton on H248 to the cytoplasm. Next a proton a transfer occurs from protonated H248 to K223 (blue arrow in state 3) converting histidine back to H248ε. MD simulation data show that protonated K223 forms an ion-pair with the anionic E140 of the Lys/Glu pair, and this destabilizes the B conformation of H248. This has an important implication because destabilization of B conformation as well as loss of hydration in the region due to loss of charge on H248 can prevent the backflow of proton on the Lys/Glu pair, thereby enhancing the rate of forward reaction.

A second proton on K353 arrives from the periplasm (state 5), resulting in higher occupancy of the A conformation of H248 (state 6), as observed in MD simulations. This is likely an energetically unstable and/or conformationally strained state because of the presence of a proton on the Lys/Glu pair and also on K299. The high energy state will relax with the loss of the proton on the Lys/Glu pair towards MrpD (state 7) and Na^+^ coupling sites. This in turn will result in pumping of Na^+^ to the periplasm, followed by the release of the proton on K299 to the cytoplasm and its reprotonation via H248 (state 8). Our MD simulation data shows that neutral K299 can adopt a proton-accepting position from protonated H248 involving conserved water molecule and S244. Once H248 donates its proton to K299, this brings the MrpA unit into its ground state, ready for the next cycle. The notion of proton release from protonated H248 to the cytoplasm via S244 is in agreement with S244A mutant data that shows no sodium tolerance, suggesting impaired proton release due to lack of hydration and altered histidine switch dynamics.

Based on the high sequence and structural similarity of MrpA subunit to ND5 subunit of complex I, as well as the above combined simulation-biochemical data, we propose a similar mechanism of proton transport in respiratory complex I, but with adaptations specific to it (see Fig. S29). Overall, we suggest that the conserved TMH8 histidine acts as a molecular switch by funneling protons between three putative proton transfer paths (Fig. 6) in both Mrp type antiporters and respiratory complex I. The unique presence of histidine in ND5/MrpA, but not in the neighboring antiporter-like subunits (ND2 or ND4/MrpD) that are closer to the coupling sites of complex I/Mrp antiporter may suggest an important role of this switching mechanism in enhancing long-range coupling.

## Methods

### Bacterial Strains, Plasmids and Mutagenesis

The bacterial strains and plasmids used in this study are listed in supplemental Table S1. Mutations were introduced by site directed mutagenesis via PCR^40^, followed by transformation in *Escherichia coli* strain XL10-Gold. The primers used in this study are listed in supplemental Table S2. For the growth and activity assays the Mrp operon was expressed from plasmids in the *E. coli* strain KNabc(DE3). In the K-12 derivate KNabc the native Na^+^/H^+^ antiporters of *E. coli* have been deleted, rendering the strain highly sensitive to the presence of sodium^29^. The gene for T7 RNA polymerase (DE3) was introduced into the KNabc chromosome by application of the “λDE3 Lysogenization Kit” (Novagen) in accordance with manufacturer specifications.

### Growth Conditions and Assays

All *E. coli* strains were cultured at 37°C. Strain XL10-Gold was grown in LB medium (pH 7.5), while strain KNabc(DE3) was grown without induction in modified LBK medium^41^ consisting of 10 g/L Tryptone, 5 g/L Yeast extract and 100 mM potassium phosphate buffer (pH 8.0). All media were supplemented with 100 μg/ml Carbenicillin.

Growth assays comparing the sodium tolerance of wild type and mutant Mrp antiporter were carried out shaking, at 180 rpm, over the course of 20 h in 96-well-plates with a culture volume of 1 ml. For this purpose, the media were supplemented with 0 – 800 mM sodium chloride. The growth was then evaluated by measuring OD600 using a SpectraMax M2/M2e Microplate Reader.

### Preparation of Everted Membrane Vesicles

Everted membrane vesicles were prepared based on a procedure described previously^42^. The harvested cells were converted to spheroplasts by treatment with 100 μg/ml lysozyme in the presence of 30 % sucrose. The spheroplasts were then suspended in TSCD buffer (10 mM Tris · HCl, 0.25 M sucrose, 140mM choline chloride, and 0.5 mM DTT; pH 7.5) and disrupted by six treatments of 30 seconds at 23 % amplitude and 60 seconds at rest in a Branson Digital Sonifier. After the removal of cell debris, the vesicles were centrifuged, resuspended in TSCD supplemented with 100 g/L glycerol and frozen in liquid nitrogen until use.

### Na^+^/H^+^ Antiport Assay

The activity of the antiporter was investigated by an assay based on the ΔpH dependent quenching of acridine orange^30^. 200 μg of everted membrane vesicles were suspended in 2 ml buffer containing 10 mM bis-tris propane · sulfate, 140 mM choline chloride, 5 mM magnesium chloride and 1 μM acridine orange (pH 9.5). The vesicles were acidified by the addition of 3.75 mM Tris · D-lactate. Antiport activity was initiated by the addition of 10 mM sodium chloride. The pH gradient was abolished by the addition of 10 mM ammonium sulfate. The measurements were taken on a Fluorolog®-3 spectrofluorometer (HORIBA Scientific) at an excitation wavelength of 492 nm and an emission wavelength of 525 nm.

### SDS-Page and Western blot

Cells were disrupted by three passes of 20 seconds at 6.0 m/s in a Savant Fastprep FP120 Cell Disruptor. Aliquots of 9 - 18 µg of total protein were separated on polyacrylamide SDS gels and then transferred to polyvinylidene difluoride (PVDF) membranes using Transfer Packs (Bio-Rad). The FLAG-tag fused to MrpG was detected by mouse Anti-DYKDDDDK Antibody (Invitrogen; RRID: AB_1957945) and the loading control was detected by mouse Anti-GAPDH Antibody (Invitrogen; RRID: AB_10977387). Goat anti-Mouse HRP (Invitrogen; RRID: AB_2533947) was used in conjunction with ECL solution (Thermo Scientific) to take chemiluminescence images on a Bio-Rad ChemiDoc imager. The GAPDH signal was used for Western blot normalization by the software Image Lab™ (BioRad; Version 6.1.0 Build 7). For comparison of expression levels, the GAPDH normalized mutant FLAG-tag signals were compared to the GAPDH normalized wild type signal of the respective Western blot. For each sample two biological replicates with three to four technical replicates each were measured (n = 7 – 8) and the results analyzed with the software Microsoft Excel.

### Computational methods

All-atom molecular dynamics (MD) simulations were conducted using the monomeric form of 2.2 Å resolution structure of Mrp antiporter from *Bacillus pseudofirmus* (PDB 7QRU^19^). The two transmembrane helices (residues 1 to 47) of chain E from the adjacent monomer were included to maintain monomer-monomer interactions at the interface. All water and lipid molecules found in the cryo-EM structure were retained for model system construction. The protonation states of amino acids were selected based on p*K*_a_ calculations on the structure of Mrp antiporter^19^ using Propka tool^43^ (see Table S3). The monomeric protein system was embedded in a hybrid lipid bilayer, consisting of a mixture of palmitoyl oleoyl phosphatidylethanolamine (POPE) and palmitoyl oleoyl phosphatidylglycerol (POPG) lipids in a ratio of 80:20. This specific lipid composition was chosen to emulate the experimental settings as close as possible^19^. The lipid bilayer was constructed using CHARMM-GUI tools^44^ and protein placement in membrane was achieved using OPM-based alignment^45^. The protein-lipid system was solvated with the TIP3P water molecules and Na^+^ and Cl^-^ ions maintaining a concentration of 150 mM. All system components were treated using the CHARMM36^46-49^ forcefield, with NBFIX^50^ corrections implemented to correct the interactions between sodium ions and protein. Wild type model systems in different protonation states as well as mutant model systems were created using VMD/PSFGEN tools^51^. The final system consisted of approximately 450000 atoms. The simulation protocol started with a (steepest descent) energy minimization on the whole system. Subsequently, an NVT equilibration at 310 K was conducted for 100 ps, employing V-rescale thermostat^52^ and applying a 2000 kJ mol^-1^ nm^-2^ harmonic restraint on all protein heavy atoms, along with the restraints on lipids phosphorus atoms, along the membrane surface. The system was then further equilibrated in NPT ensemble (maintaining 310 K temperature and 1 atm pressure) in two steps: first, a 1 ns equilibration using V-rescale thermostat^52^ and Berendsen barostat^53^, with constraints applied solely to protein backbone atoms, then a 10 ns equilibration without any restraints. Production runs were conducted utilizing the Nosé-Hoover thermostat^54,55^ and Parrinello-Rahman barostat^56^. A simulation timestep of 2 fs was achieved with the LINCS algorithm^57^. The electrostatic interactions were computed with the particle-mesh Ewald (PME) method^58^ with a cutoff of 12 Å.

All production MD simulations are described in Table S4. Each system was replicated three times, with each simulation replica originating from the same initial configuration but independently minimized and equilibrated. All MD simulations and trajectory processing were carried out using GROMACS 2021.5^59^. Trajectory visual analysis and volumetric maps calculations were performed using Visual Molecular Dynamics (VMD) software^51^. For data analysis, MDtraj software was used^60^, and figures were generated with VMD, Pymol, and Inkscape. The secondary structure analysis was performed with the DSSP (Define Secondary Structure of Proteins) tool^61,62^ as implemented in MDTraj. Trajectory frames were extracted at a regular interval of 1 ns, and each frame is processed using MDTraj/DSSP to assign a secondary structure label to every residue of the H248 helix.

## Acknowledgements

VS acknowledges research funding from the Research Council of Finland, the Jane and Aatos Erkko Foundation, the Sigrid Jusélius Foundation, the Magnus Ehrnrooth Foundation, the Cancer Foundation and the University of Helsinki. VZ acknowledges funding by the Deutsche Forschugsgemeinschaft (German Research Foundation), grant CRC1507/P14. The Center for Scientific Computing (CSC), Finland is acknowledged for excellent high-performance computing resources, as well as the IT support by the Faculty of Science, University of Helsinki. We thank Karin Siegmund for excellent technical support.

The authors declare that they have no conflict of interest.

## Notes

### Competing Interest Statement

The authors have declared no competing interest.

