## Supplementary Data for "Conformational dynamics of a histidine molecular switch in a cation/proton antiporter"

**Figure/table contents**

| Figure/Table | Content |
|---|---|
| S1 | SDS-PAGE |
| S2 | WT, simulation systems 1, 2 |
| S3 | T306V, simulation systems 30, 31 |
| S4 | S146A, simulation systems 6, 7 |
| S5 | S146T, simulation systems 8, 9 |
| S6 | L247H, simulation systems 14, 15 |
| S7 | L247H, simulation systems 16, 17 |
| S8 | S249H, simulation systems 22, 23 |
| S9 | S249H, simulation systems 24, 25 |
| S10 | S244A, simulation systems 12, 13 |
| S11 | H248A, H248F, simulation systems 18, 19 |
| S12 | T251V, simulation systems 28, 29 |
| S13 | F119L, simulation systems 4, 5 |
| S14 | A250V, simulation systems 26, 27 |
| S15 | W232A, simulation systems 10, 11 |
| S16 | S249A, simulation systems 20, 21 |
| S17 | WT, simulation systems 34, 35 |
| S18 | WT, simulation systems 36, 37 |
| S19 | WT, simulation systems 43, 44 |
| S20 | WT, simulation systems 32, 33 |
| S21 | WT, simulation systems 38, 39 |
| S22 | WT, simulation systems 40, 41 |
| S23 | WT, simulation systems 3, 42 |
| S24 | WT, simulation system 45 |
| S25 | WT, simulation systems 46, 47 |
| S26 | W232A, H248A and H248F mutations and intermediate position of H248 |
| S27 | Helicity of H248 carrying segment |
| S28 | Hydration, conformational and protonation state changes |
| S29 | Histidine switch mechanism in complex I |
| S30 | Conformational occupancies |
| S31 | Conformational occupancies |
| S32 | L247H and S249H mutations |
| S33 | S146A, T306V, simulation systems 48,49. Conformational occupancies |
| Table S1 | Strains and plasmids |
| Table S2 | Primers |
| Table S3 | Predicted protonation states |
| Table S4 | Simulation systems |

**Figure S1 Expression level of wild type and mutant BpMrp in KNabc(DE3).** Detection of FLAG-tagged MrpG and GAPDH loading control in whole cell extracts from *Escherichia coli* KNabc(DE3). The expression of BpMrp wild type, negative control (plasmid pUB26) or BpMrp mutants was analyzed by immunoblotting following SDS-PAGE. Additionally, purified wild type Mrp antiporter was utilized as positive control. For each sample, three to four technical replicates of two separate preparations were examined (n = 7 – 8). Example blots are shown in **(a)** and **(b)**. For each blot the obtained chemiluminescence intensities of the FLAG-MrpG bands were normalized to the GAPDH loading control and then the mutants were compared to the wild type. The resulting distribution of apparent expression levels is shown in form of a boxplot in **(c)**. The X marks the mean value and the horizontal line the median value.

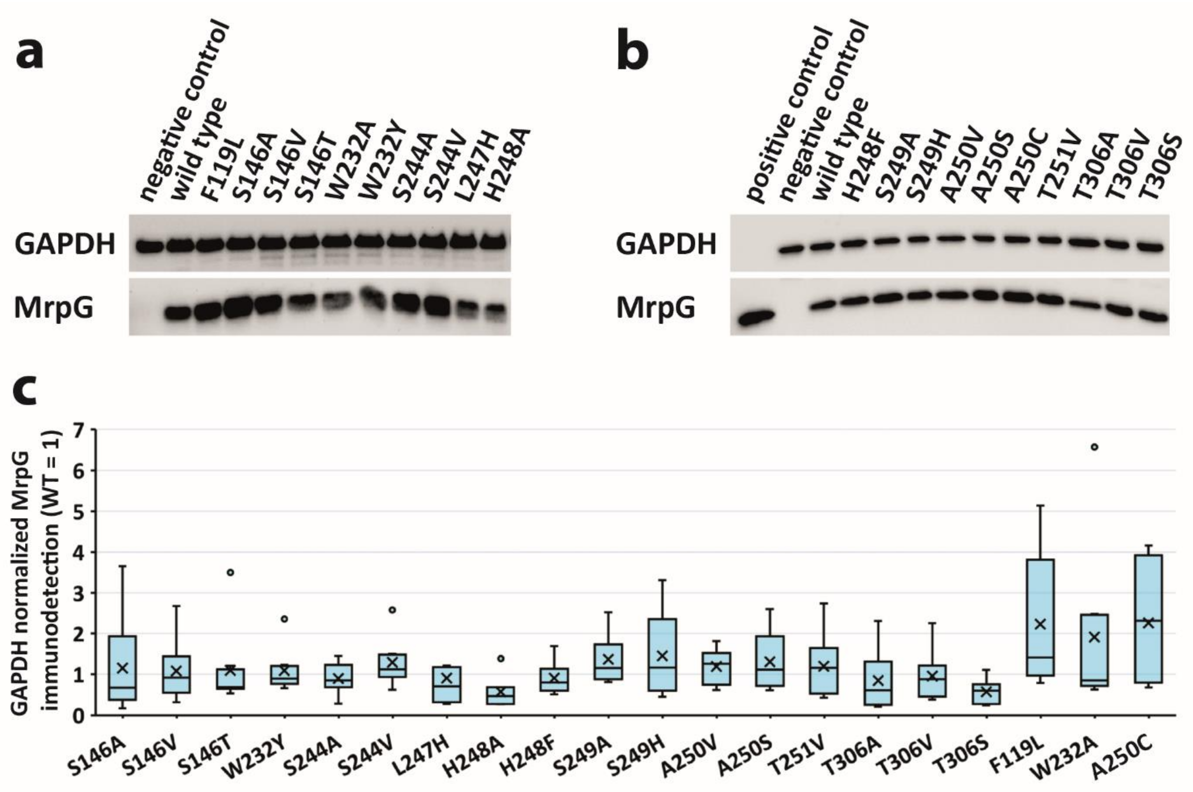

**Figure S2. Dynamics of histidine sidechain and backbone carrying it.** Data from wild type simulation setups number 1 and 2. **(a)** Scatter plots display distances of S146 and T306 side chains from H248 side chain. Data is shown from all three simulation replicas in three different colors. Left – H248δ and Right – H248ε **(b)** Time resolved secondary structure analysis of the H248 carrying segment A245-A255. The three rows represent the different simulation replicas. Left – H248δ and Right – H248ε. See main text Fig. 3 for more details.

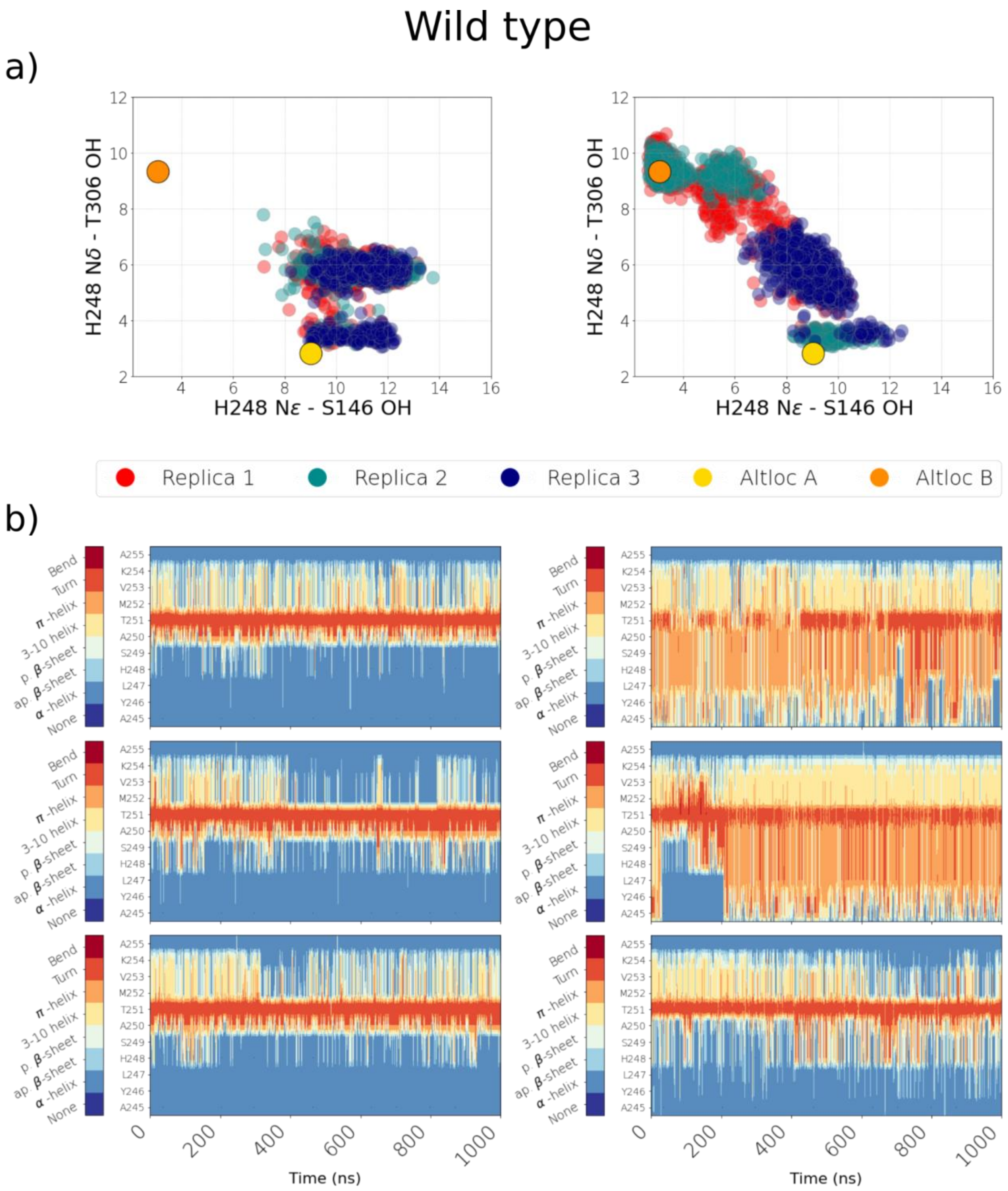

**Figure S3. Dynamics of histidine sidechain and backbone carrying it.** Data from T306V simulation setups number 30 and 31. **(a)** Scatter plots display distances of S146 and T306 side chains from H248 side chain. Data is shown from all three simulation replicas in three different colors. Left – H248δ and Right – H248ε **(b)** Time resolved secondary structure analysis of the H248 carrying segment A245-A255. The three rows represent three different simulation replicas. Left – H248δ and Right – H248ε. See main text Fig. 3 for more details.

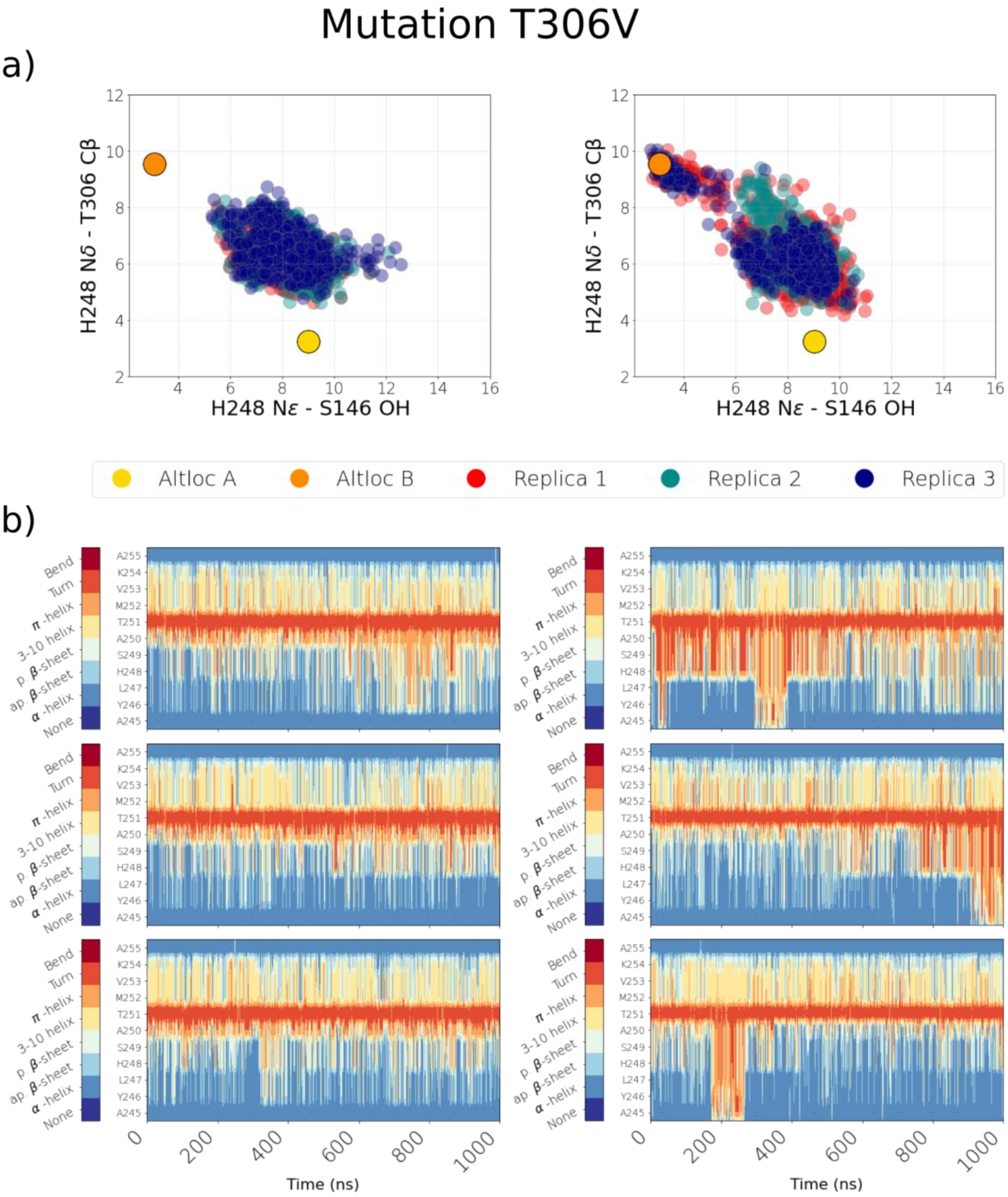

**Figure S4. Dynamics of histidine sidechain and backbone carrying it.** Data from S146A simulation setups number 6 and 7. **(a)** Scatter plots display distances of S146 and T306 side chains from H248 side chain. Data is shown from all three simulation replicas in three different colors. Left – H248δ and Right – H248ε **(b)** Time resolved secondary structure analysis of the H248 carrying segment A245-A255. The three rows represent three different simulation replicas. Left – H248δ and Right – H248ε. See main text Fig. 3 for more details.

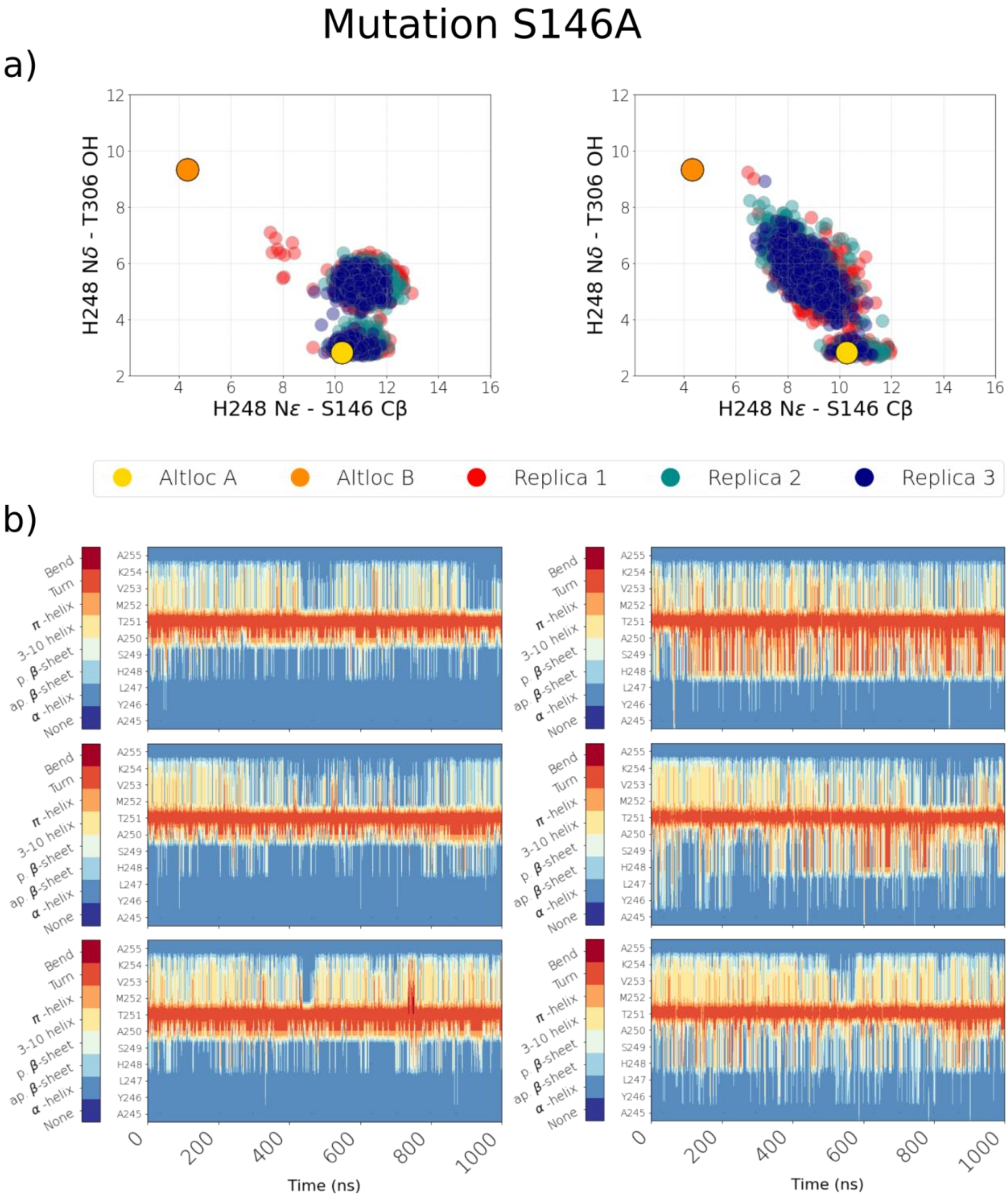

**Figure S5. Dynamics of histidine sidechain and backbone carrying it.** Data from S146T simulation setups 8 and 9. **(a)** Scatter plots display distances of S146 and T306 side chains from H248 side chain. Data is shown from all three simulation replicas in three different colors. Left – H248δ and Right – H248ε **(b)** Time resolved secondary structure analysis of the H248 carrying segment A245-A255. The three rows represent three different simulation replicas. Left – H248δ and Right – H248ε. See main text Fig. 3 for more details.

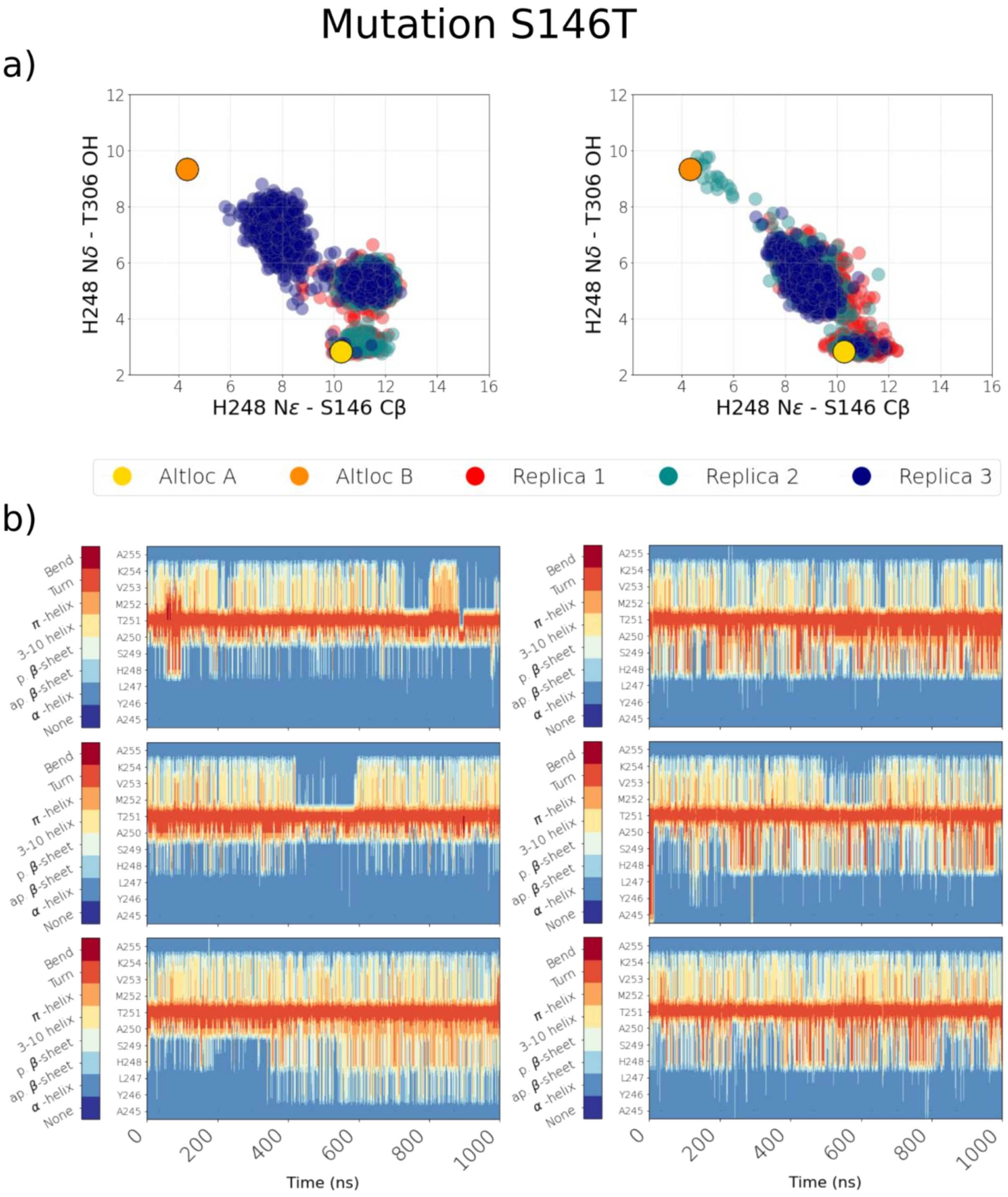

**Figure S6. Dynamics of histidine sidechain and backbone carrying it.** Data from L247H simulation setups number 14 and 15. **(a)** Scatter plots display distances of S146 and T306 side chains from H248 side chain. Data is shown from all three simulation replicas in three different colors. Left – H248δ and Right – H248ε **(b)** Time resolved secondary structure analysis of the H248 carrying segment A245-A255. The three rows represent three different simulation replicas. Left – H248δ and Right – H248ε. See main text Fig. 3 for more details.

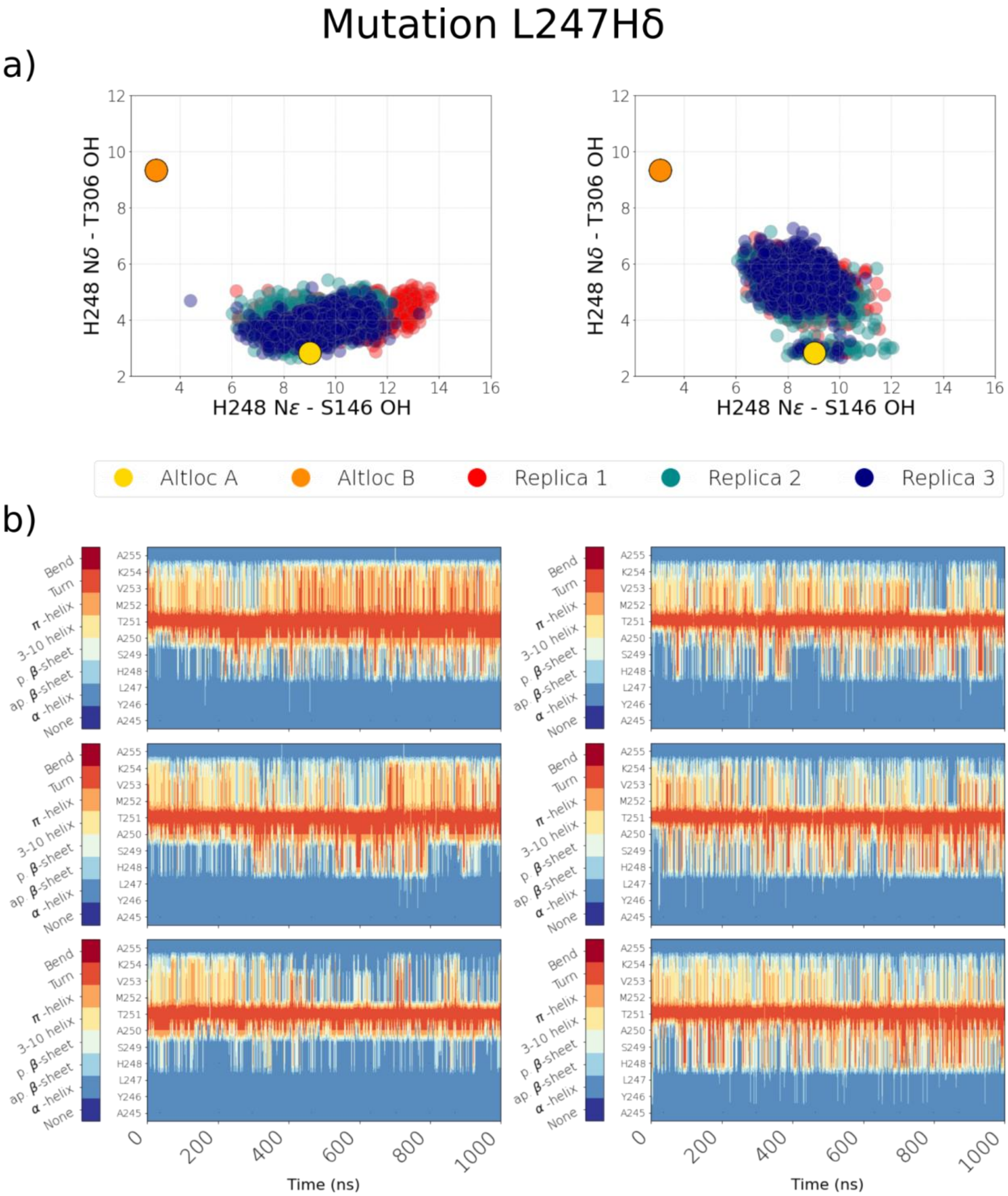

**Figure S7. Dynamics of histidine sidechain and backbone carrying it.** Data from L247H simulation setups number 16 and 17. **(a)** Scatter plots display distances of S146 and T306 side chains from H248 side chain. Data is shown from all three simulation replicas in three different colors. Left – H248δ and Right – H248ε **(b)** Time resolved secondary structure analysis of the H248 carrying segment A245-A255. The three rows represent three different simulation replicas. Left – H248δ and Right – H248ε. See main text Fig. 3 for more details.

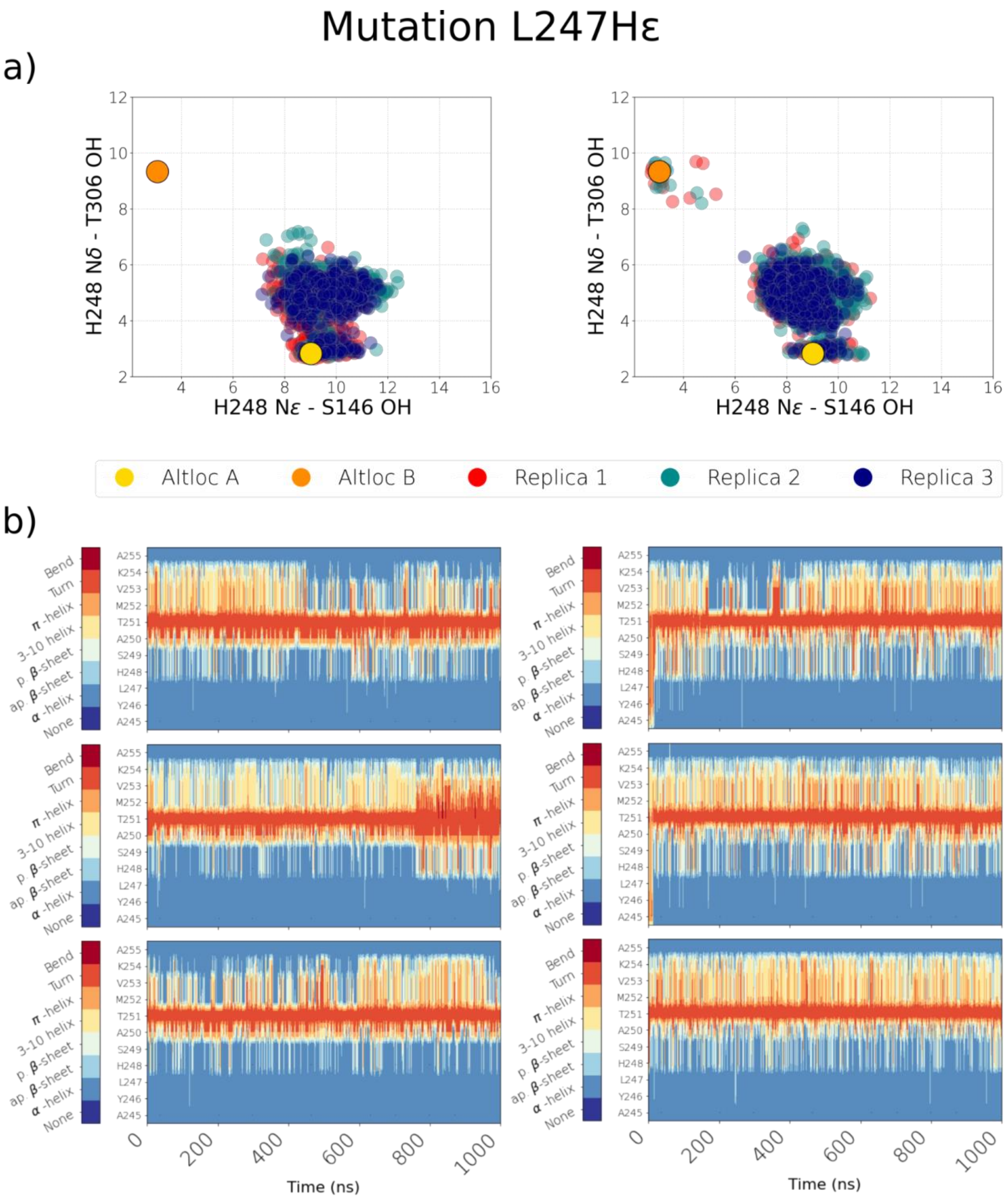

**Figure S8. Dynamics of histidine sidechain and backbone carrying it.** Data from S249H simulation setups number 22 and 23. **(a)** Scatter plots display distances of S146 and T306 side chains from H248 side chain. Data is shown from all three simulation replicas in three different colors. Left – H248δ and Right – H248ε **(b)** Time resolved secondary structure analysis of the H248 carrying segment A245-A255. The three rows represent three different simulation replicas. Left – H248δ and Right – H248ε. See main text Fig. 3 for more details.

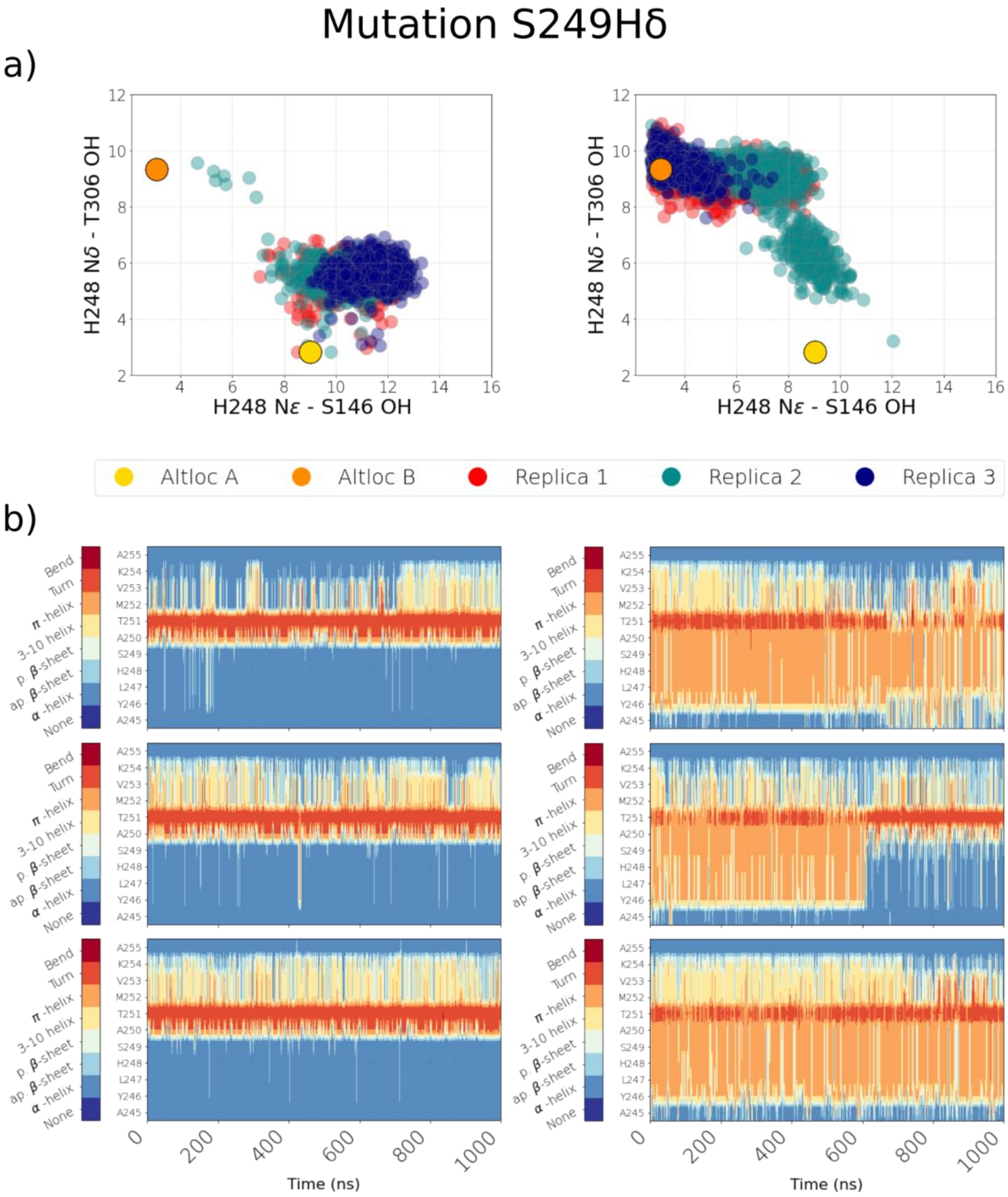

**Figure S9. Dynamics of histidine sidechain and backbone carrying it.** Data from S249H simulation setups number 24 and 25. **(a)** Scatter plots display distances of S146 and T306 side chains from H248 side chain. Data is shown from all three simulation replicas in three different colors. Left – H248δ and Right – H248ε **(b)** Time resolved secondary structure analysis of the H248 carrying segment A245-A255. The three rows represent three different simulation replicas. Left – H248δ and Right – H248ε. See main text Fig. 3 for more details.

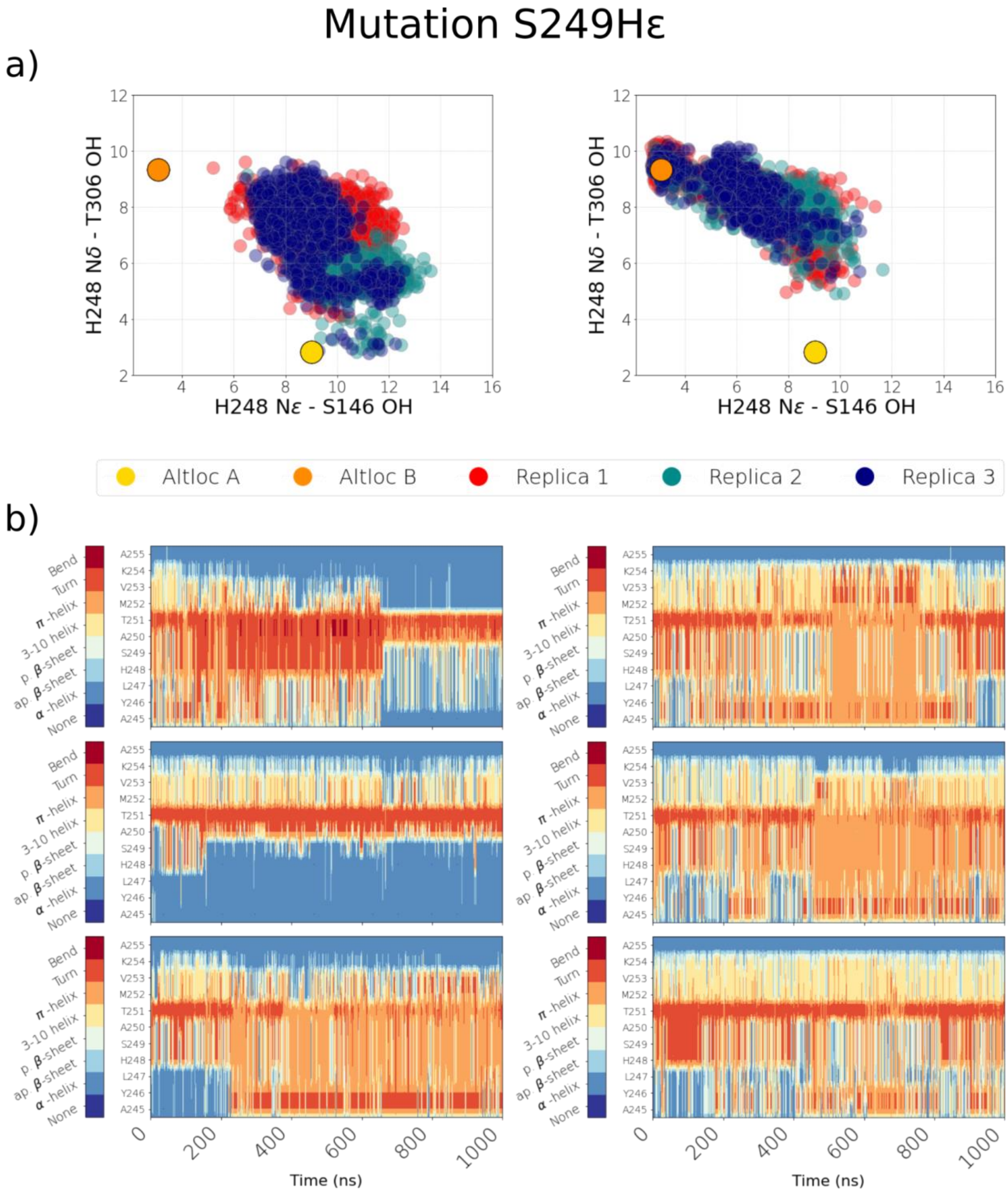

**Figure S10. Dynamics of histidine sidechain and backbone carrying it.** Data from S244A simulation setups number 12 and 13. **(a)** Scatter plots display distances of S146 and T306 side chains from H248 side chain. Data is shown from all three simulation replicas in three different colors. Left – H248δ and Right – H248ε **(b)** Time resolved secondary structure analysis of the H248 carrying segment A245-A255. The three rows represent three different simulation replicas. Left – H248δ and Right – H248ε. See main text Fig. 3 for more details.

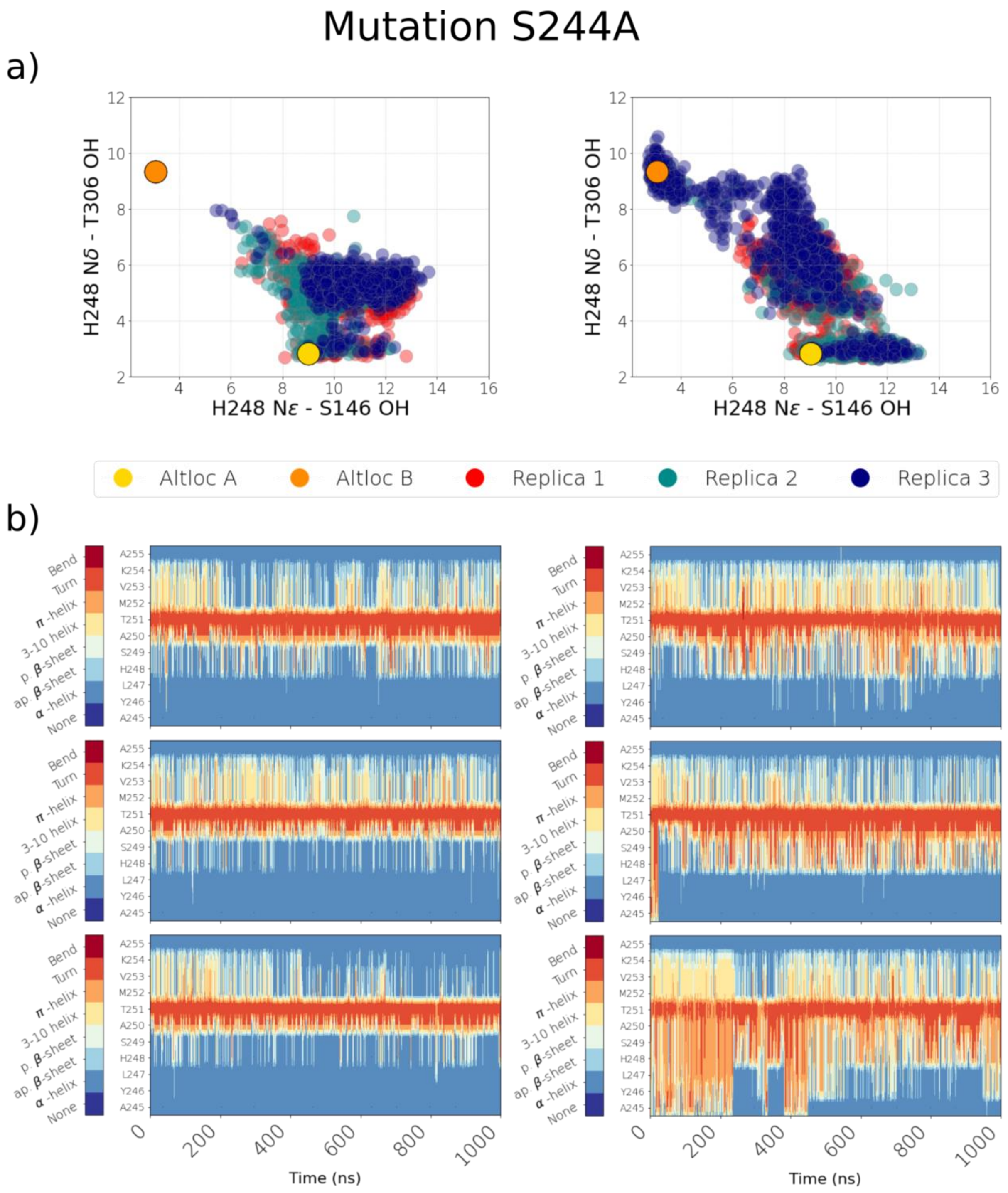

**Figure S11. Dynamics of histidine sidechain and backbone carrying it.** Data from H248A **(a)** and H248F **(b)** simulation setups number 18 and 19. Time resolved secondary structure analysis of the H248 carrying segment A245-A255. The three rows represent three different simulation replicas.

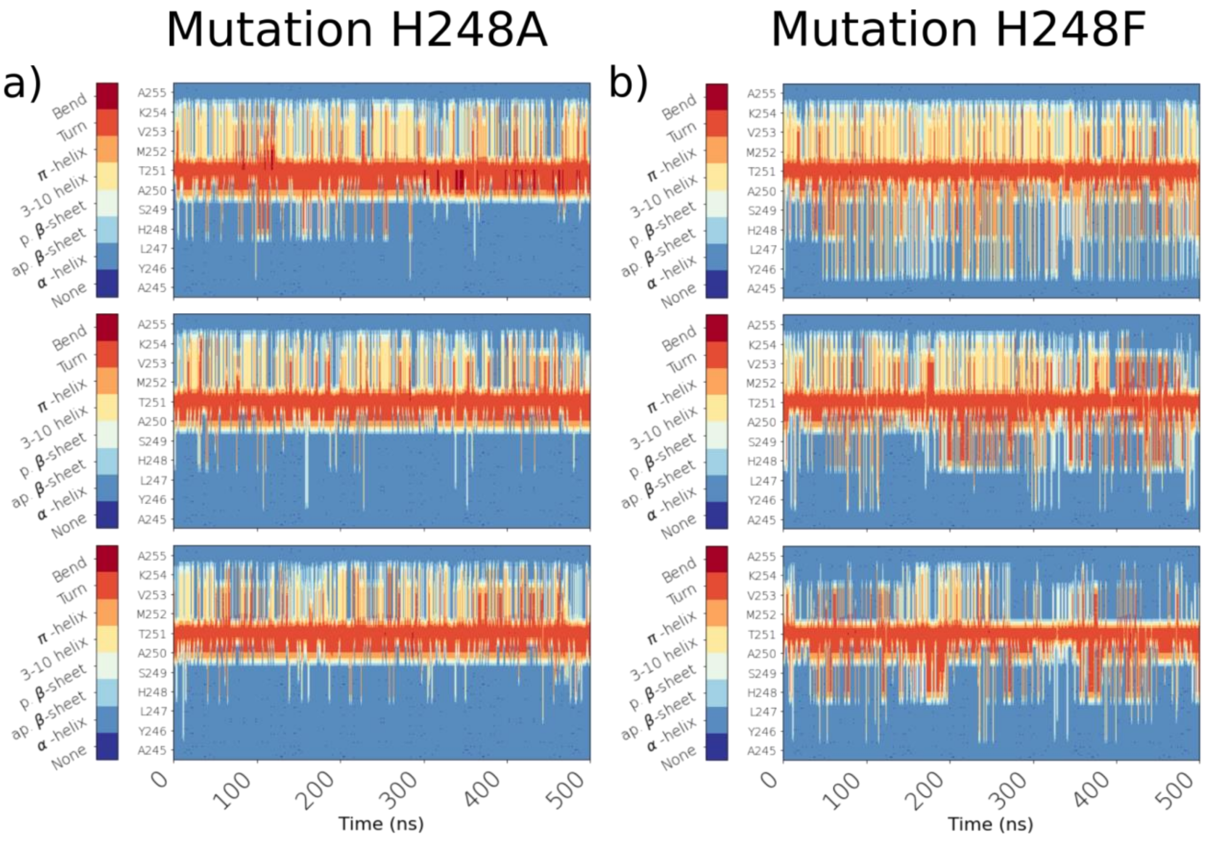

**Figure S12. Dynamics of histidine sidechain and backbone carrying it.** Data from T251V simulation setups number 28 and 29. **(a)** Scatter plots display distances of S146 and T306 side chains from H248 side chain. Data is shown from all three simulation replicas in three different colors. Left – H248δ and Right – H248ε **(b)** Time resolved secondary structure analysis of the H248 carrying segment A245-A255. The three rows represent three different simulation replicas. Left – H248δ and Right – H248ε. See main text Fig. 3 for more details.

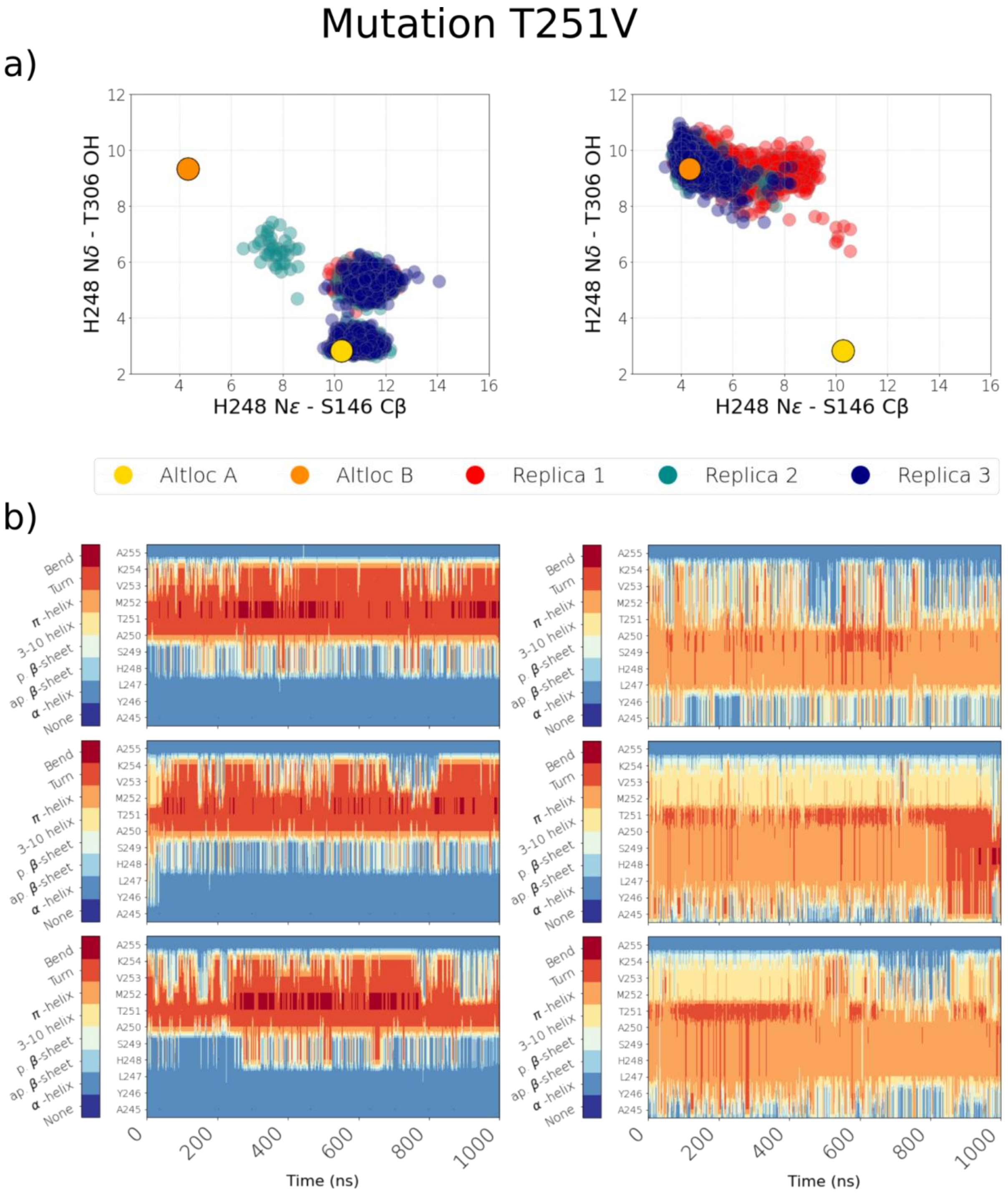

**Figure S13. Dynamics of histidine sidechain and backbone carrying it.** Data from F119L simulation setups number 4 and 5. **(a)** Scatter plots display distances of S146 and T306 side chains from H248 side chain. Data is shown from all three simulation replicas in three different colors. Left – H248δ and Right – H248ε **(b)** Time resolved secondary structure analysis of the H248 carrying segment A245-A255. The three rows represent three different simulation replicas. Left – H248δ and Right – H248ε. See main text Fig. 3 for more details.

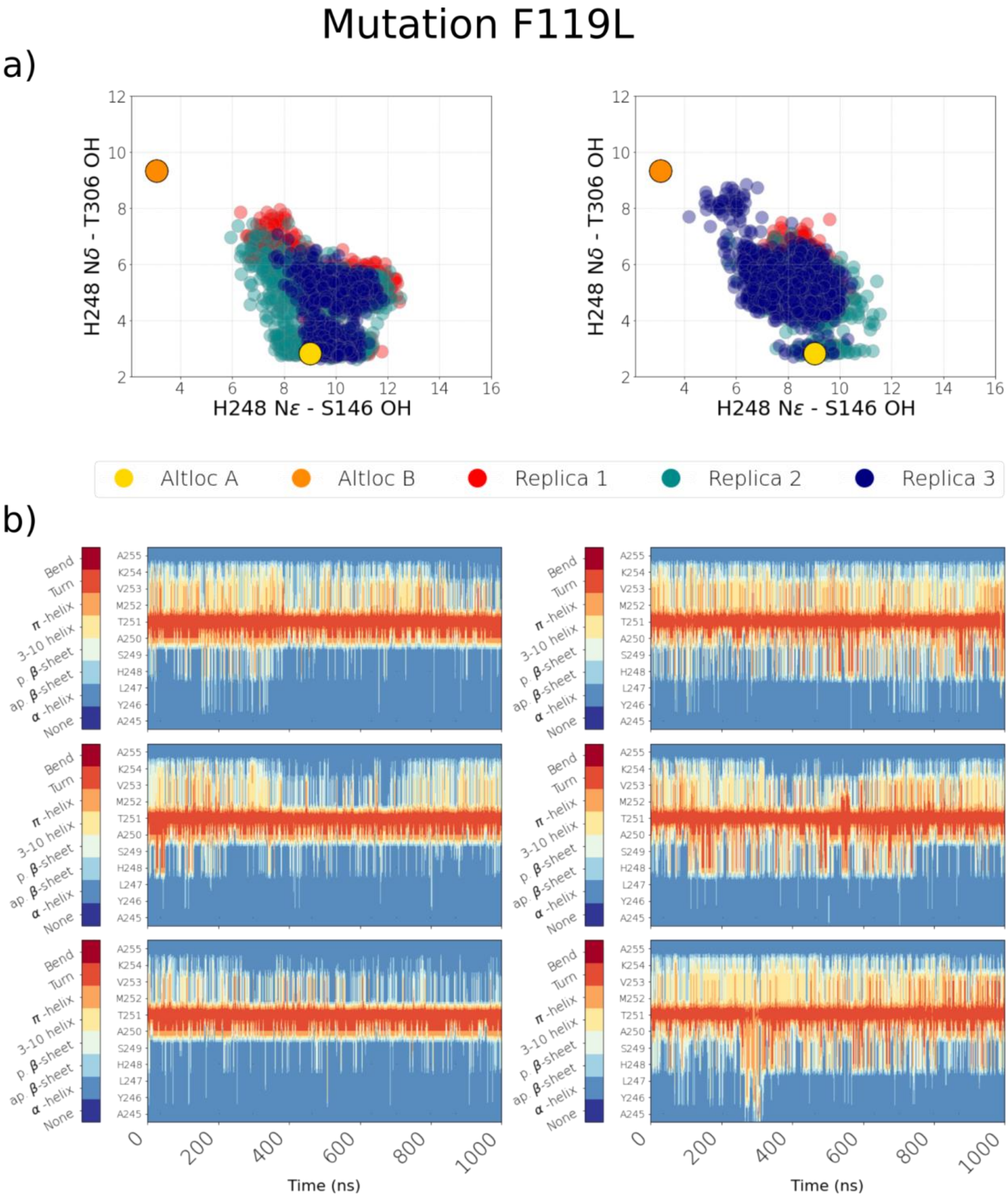

**Figure S14. Dynamics of histidine sidechain and backbone carrying it.** Data from A250V simulation setups number 26 and 27. **(a)** Scatter plots display distances of S146 and T306 side chains from H248 side chain. Data is shown from all three simulation replicas in three different colors. Left – H248δ and Right – H248ε **(b)** Time resolved secondary structure analysis of the H248 carrying segment A245-A255. The three rows represent three different simulation replicas. Left – H248δ and Right – H248ε. See main text Fig. 3 for more details.

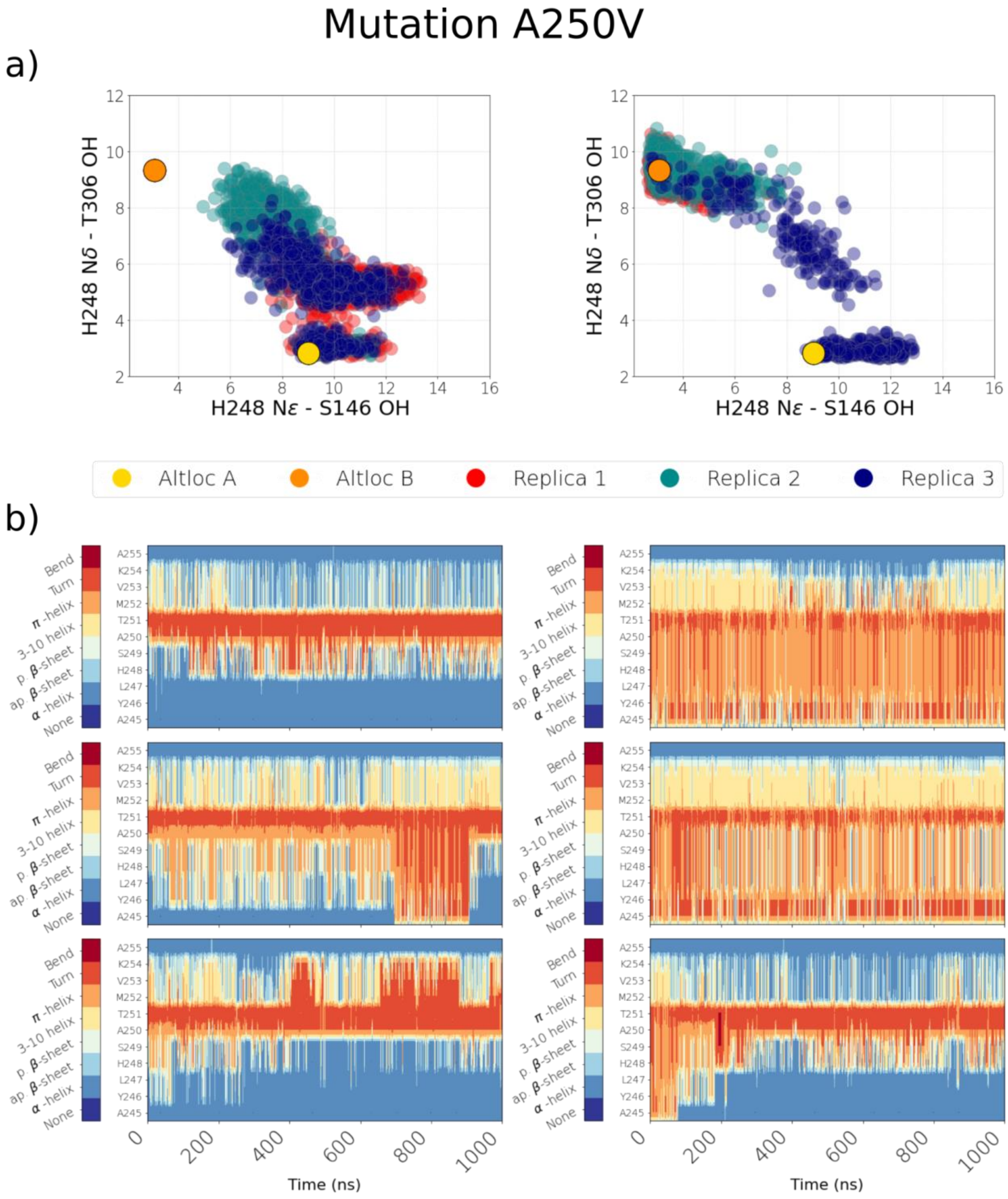

**Figure S15. Dynamics of histidine sidechain and backbone carrying it.** Data from W232A simulation setups number 10 and 11. **(a)** Scatter plots display distances of S146 and T306 side chains from H248 side chain. Data is shown from all three simulation replicas in three different colors. Left – H248δ and Right – H248ε **(b)** Time resolved secondary structure analysis of the H248 carrying segment A245-A255. The three rows represent three different simulation replicas. Left – H248δ and Right – H248ε. See main text Fig. 3 for more details.

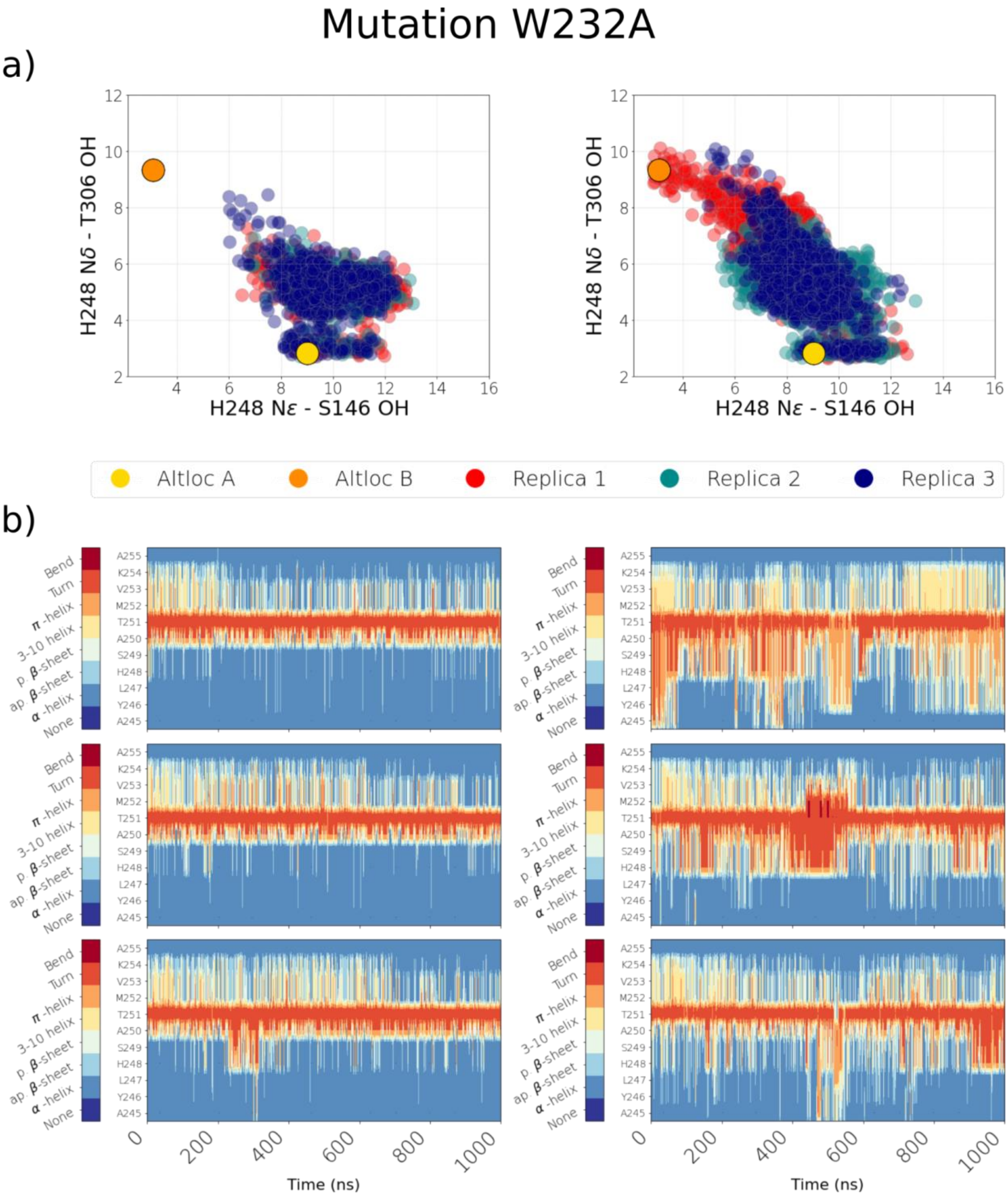

**Figure S16. Dynamics of histidine sidechain and backbone carrying it.** Data from S249A simulation setups number 20 and 21. **(a)** Scatter plots display distances of S146 and T306 side chains from H248 side chain. Data is shown from all three simulation replicas in three different colors. Left – H248δ and Right – H248ε **(b)** Time resolved secondary structure analysis of the H248 carrying segment A245-A255. The three rows represent three different simulation replicas. Left – H248δ and Right – H248ε. See main text Fig. 3 for more details.

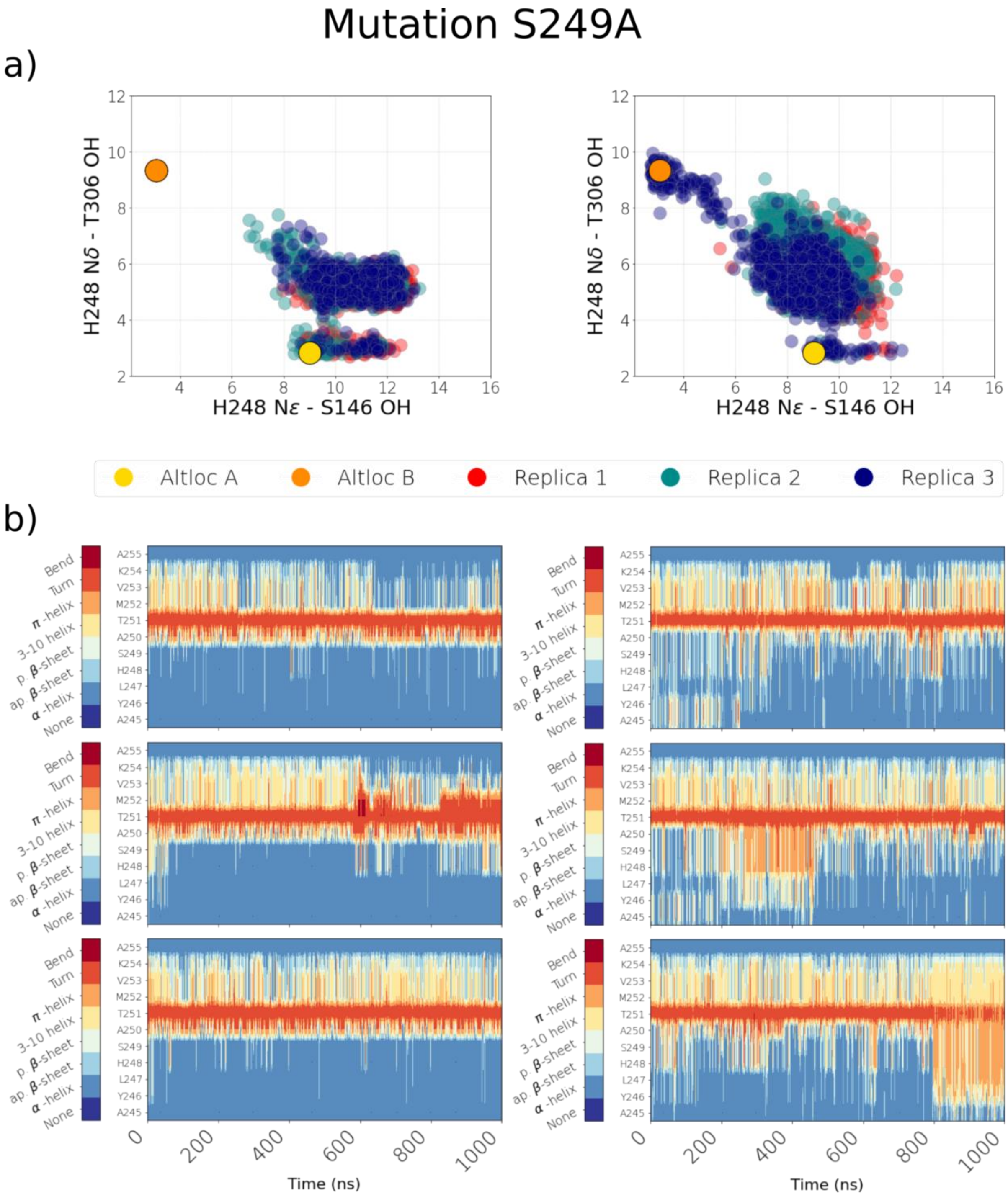

**Figure S17. Dynamics of histidine sidechain and backbone carrying it.** Data from wild type simulation setups number 34 and 35. **(a)** Scatter plots display distances of S146 and T306 side chains from H248 side chain. Data is shown from all three simulation replicas in three different colors. Left – H248δ and Right – H248ε **(b)** Time resolved secondary structure analysis of the H248 carrying segment A245-A255. The three rows represent three different simulation replicas. Left – H248δ and Right – H248ε. See main text Fig. 3 for more details.

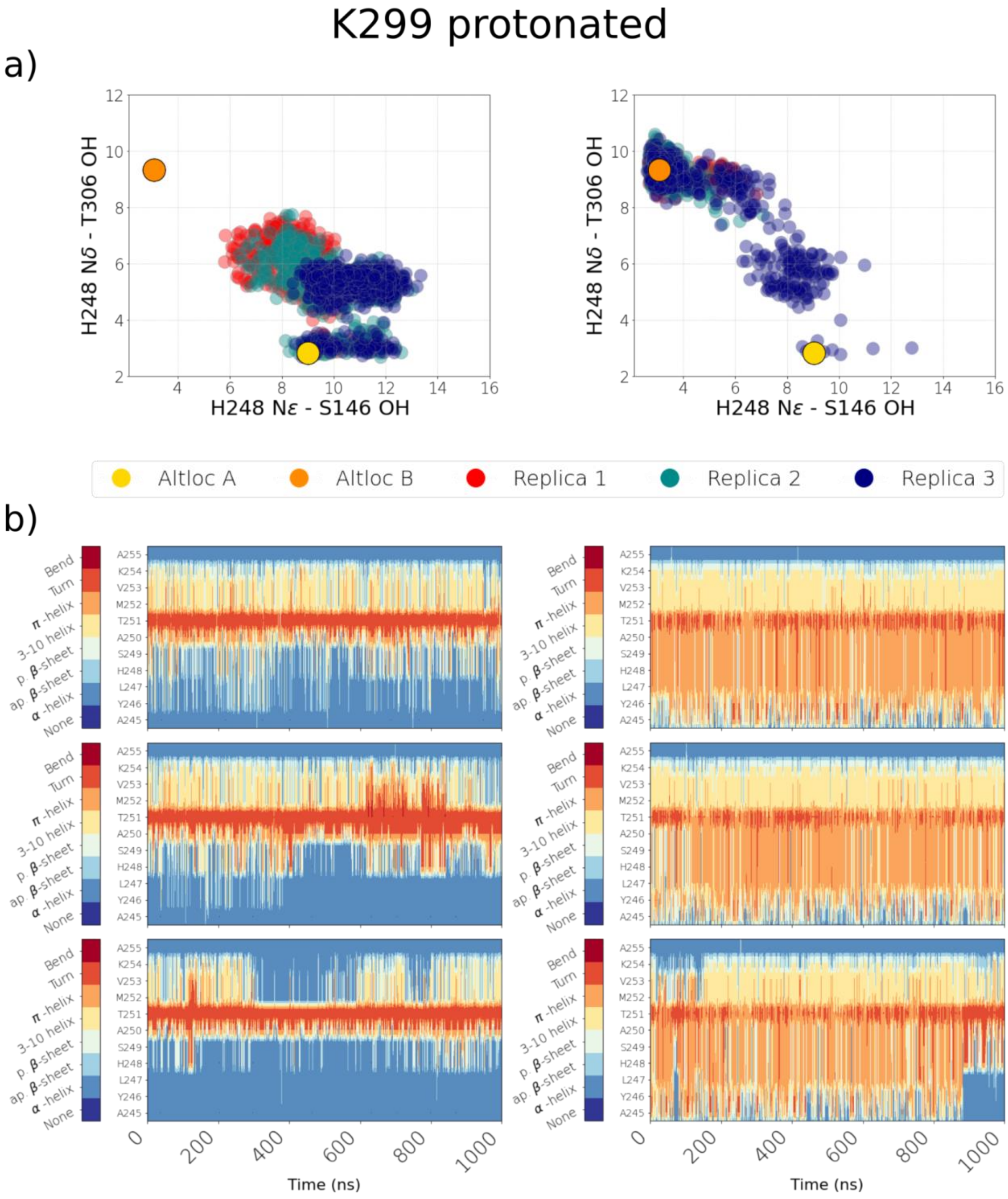

**Figure S18. Dynamics of histidine sidechain and backbone carrying it.** Data from wild type simulation setups number 36 and 37. **(a)** Scatter plots display distances of S146 and T306 side chains from H248 side chain. Data is shown from all three simulation replicas in three different colors. Left – H248δ and Right – H248ε **(b)** Time resolved secondary structure analysis of the H248 carrying segment A245-A255. The three rows represent three different simulation replicas. Left – H248δ and Right – H248ε. See main text Fig. 3 for more details.

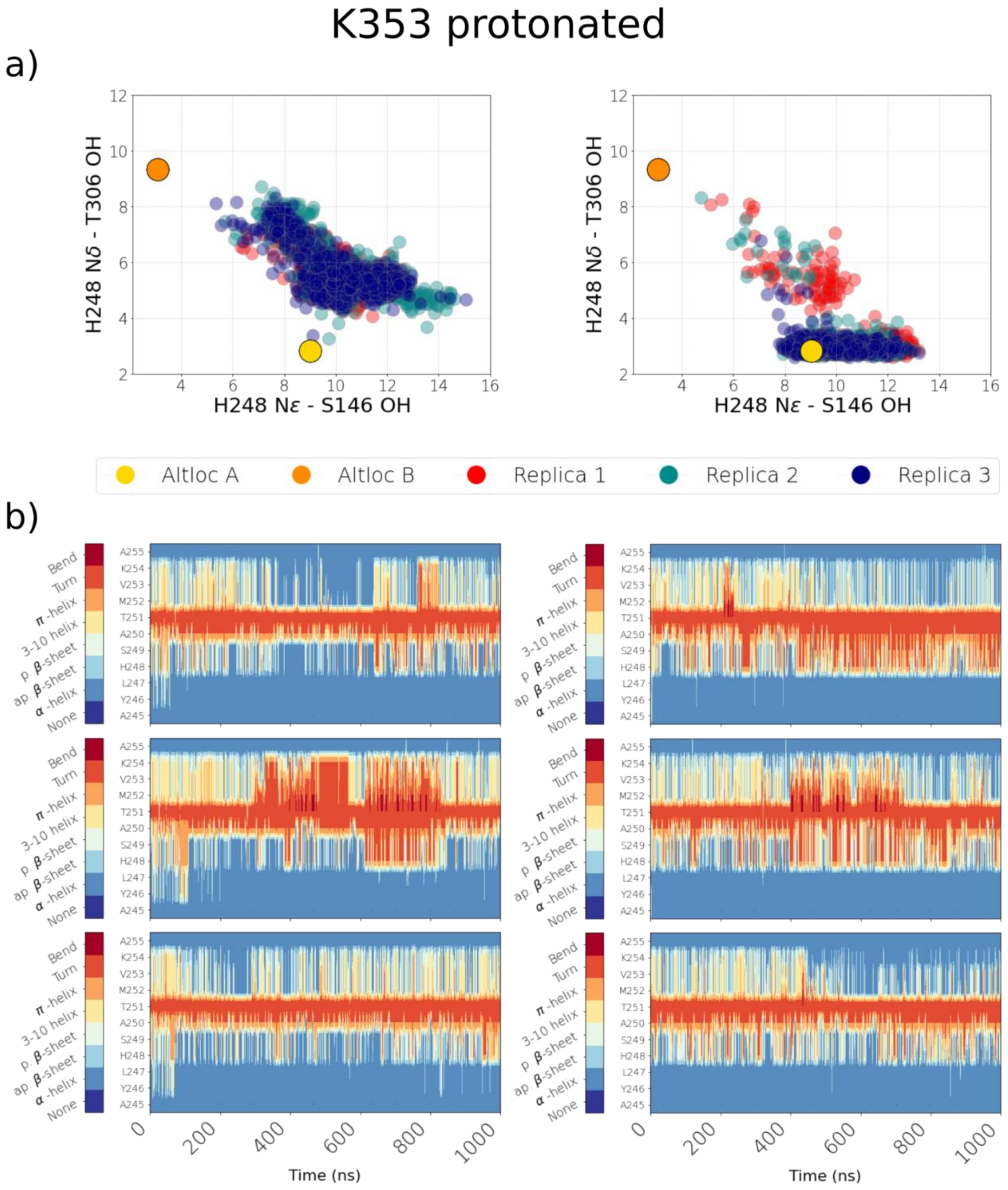

**Figure S19. Dynamics of histidine sidechain and backbone carrying it.** Data from wild type simulation setups number 43 and 44. **(a)** Scatter plots display distances of S146 and T306 side chains from H248 side chain. Data is shown from all three simulation replicas in three different colors. Left – H248δ and Right – H248ε **(b)** Time resolved secondary structure analysis of the H248 carrying segment A245-A255. The three rows represent three different simulation replicas. Left – H248δ and Right – H248ε. See main text Fig. 3 for more details.

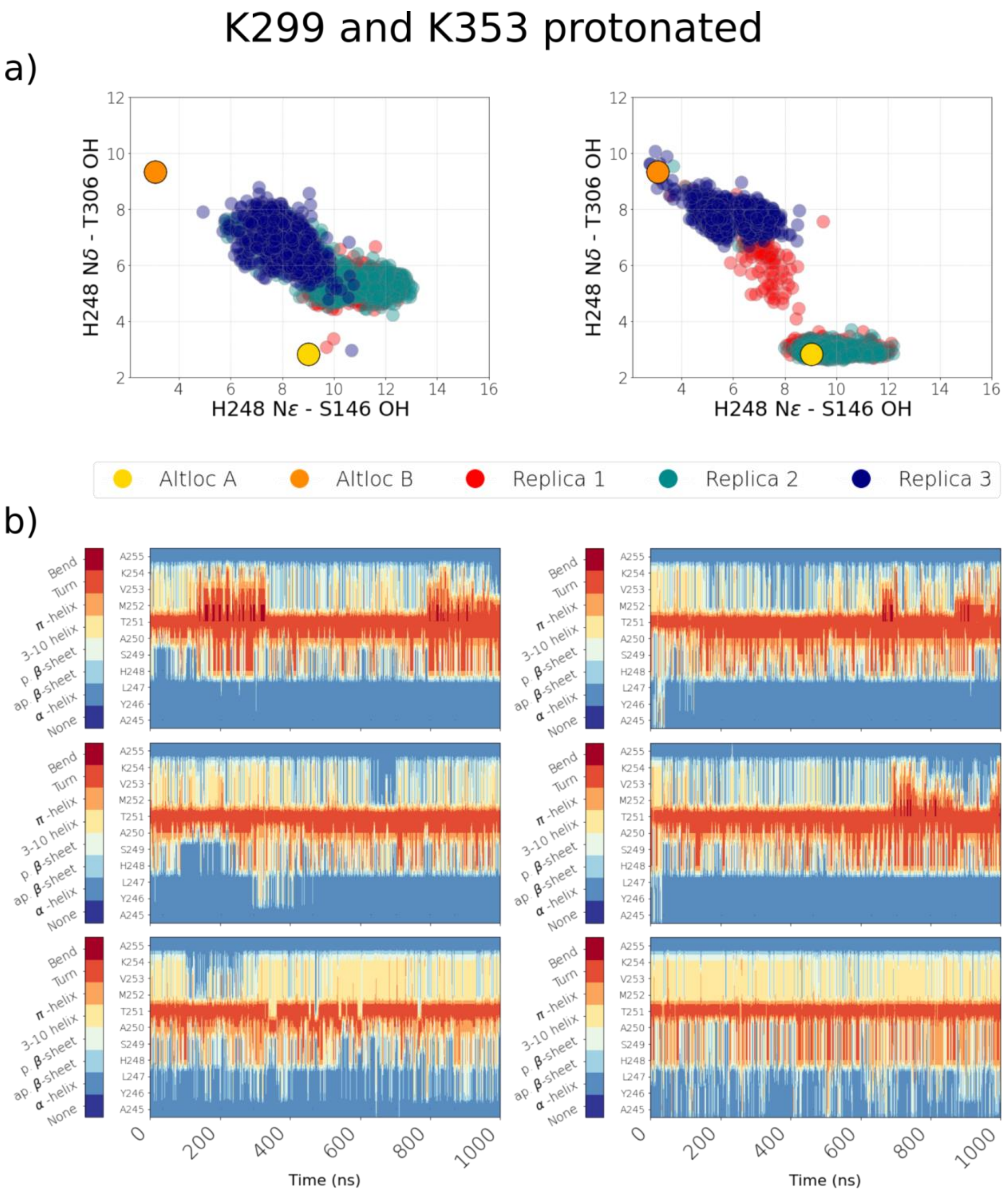

**Figure S20. Dynamics of histidine sidechain and backbone carrying it.** Data from wild type simulation setups number 32 and 33. **(a)** Scatter plots display distances of S146 and T306 side chains from H248 side chain. Data is shown from all three simulation replicas in three different colors. Left – H248δ and Right – H248ε **(b)** Time resolved secondary structure analysis of the H248 carrying segment A245-A255. The three rows represent three different simulation replicas. Left – H248δ and Right – H248ε. See main text Fig. 3 for more details.

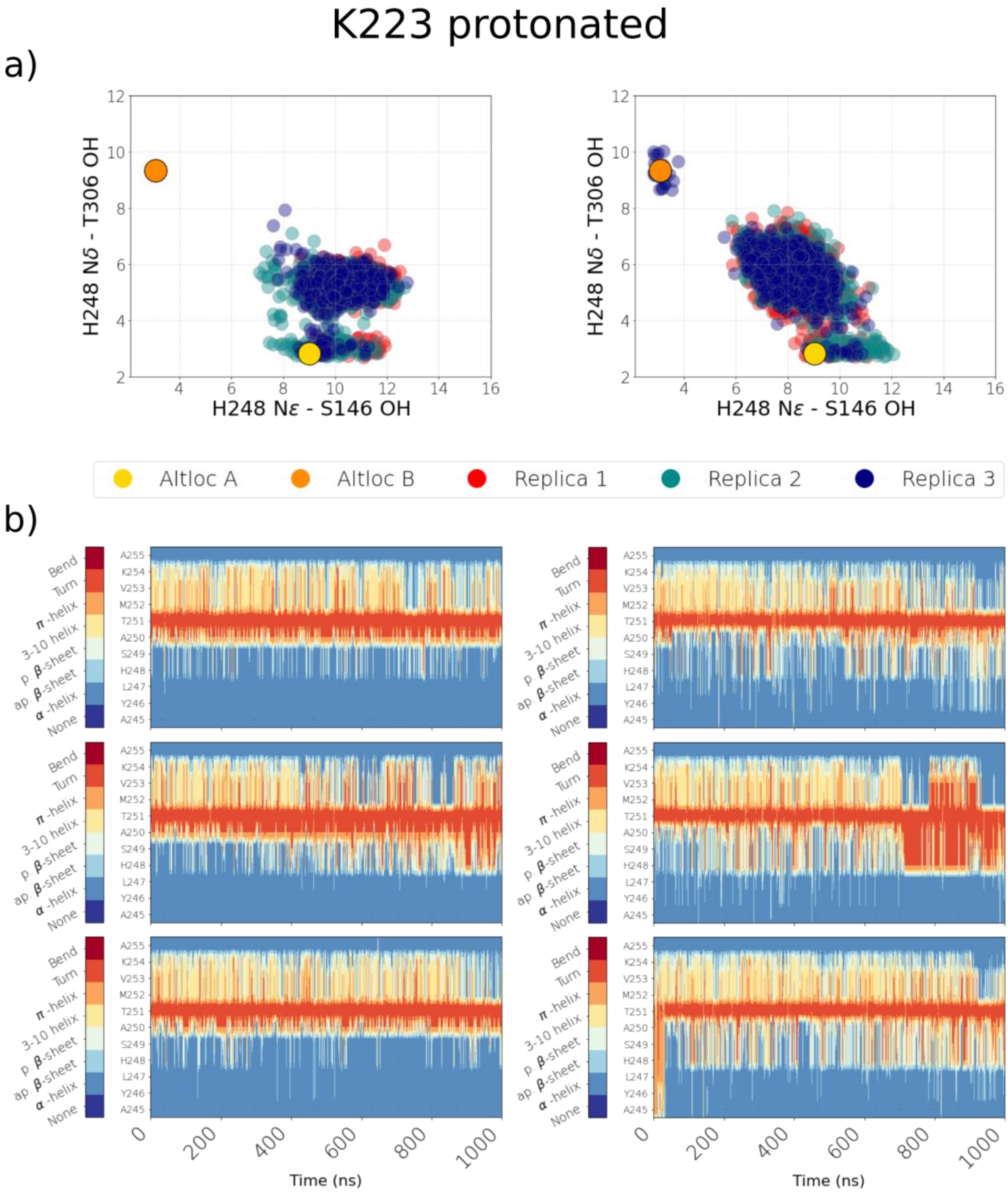

**Figure S21. Dynamics of histidine sidechain and backbone carrying it.** Data from wild type simulation setups number 38 and 39. **(a)** Scatter plots display distances of S146 and T306 side chains from H248 side chain. Data is shown from all three simulation replicas in three different colors. Left – H248δ and Right – H248ε **(b)** Time resolved secondary structure analysis of the H248 carrying segment A245-A255. The three rows represent three different simulation replicas. Left – H248δ and Right – H248ε. See main text Fig. 3 for more details.

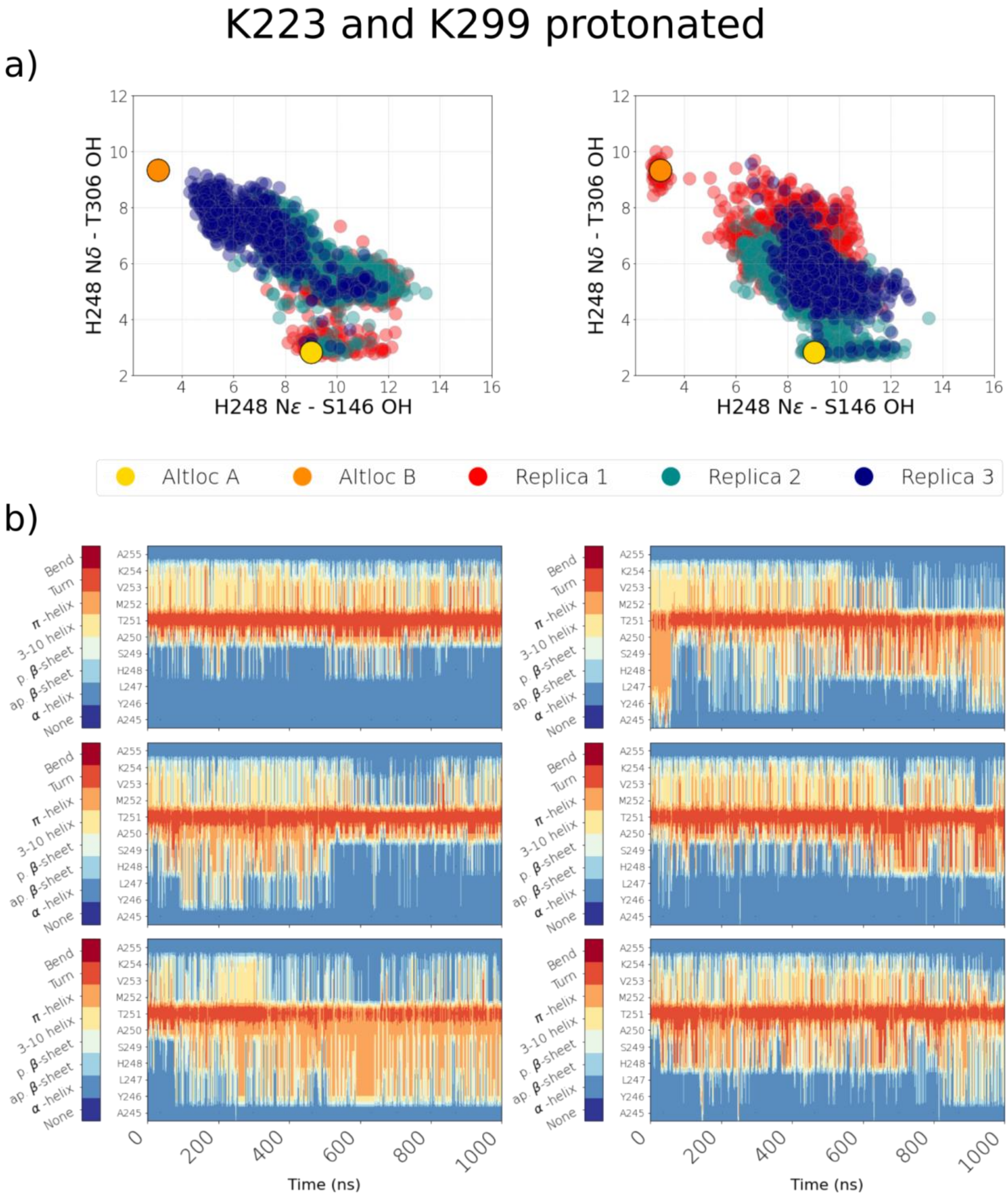

**Figure S22. Dynamics of histidine sidechain and backbone carrying it.** Data from wild type simulation setups number 40 and 41. **(a)** Scatter plots display distances of S146 and T306 side chains from H248 side chain. Data is shown from all three simulation replicas in three different colors. Left – H248δ and Right – H248ε **(b)** Time resolved secondary structure analysis of the H248 carrying segment A245-A255. The three rows represent three different simulation replicas. Left – H248δ and Right – H248ε. See main text Fig. 3 for more details.

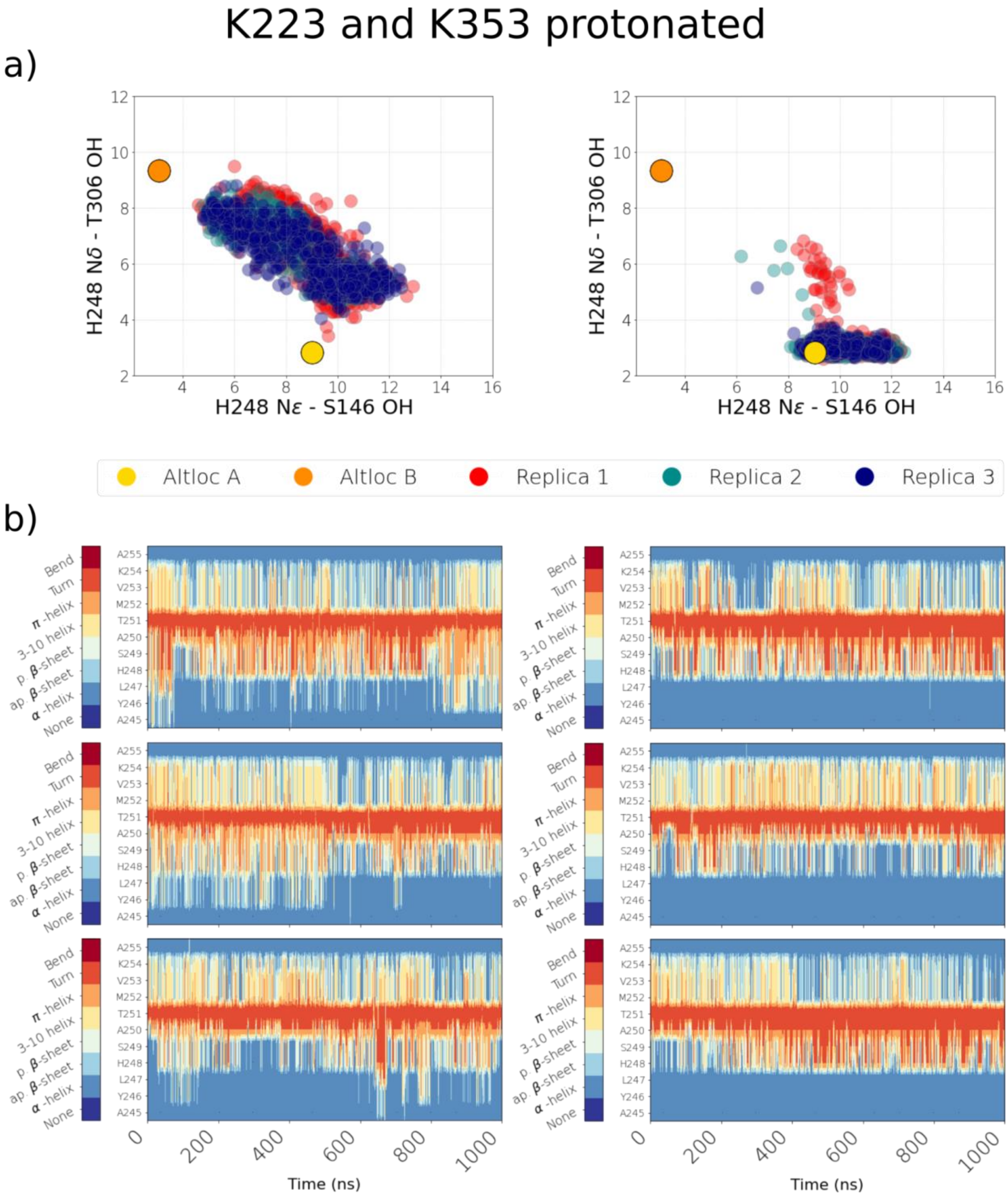

**Figure S23. Dynamics of histidine sidechain and backbone carrying it.** Data from wild type simulation setups number 3 and 42. **(a)** Scatter plots display distances of S146 and T306 side chains from H248 side chain. Data is shown from all three simulation replicas in three different colors. **(b)** Time resolved secondary structure analysis of the H248 carrying segment A245-A255. The three rows represent three different simulation replicas. See main text Fig. 3 for more details.

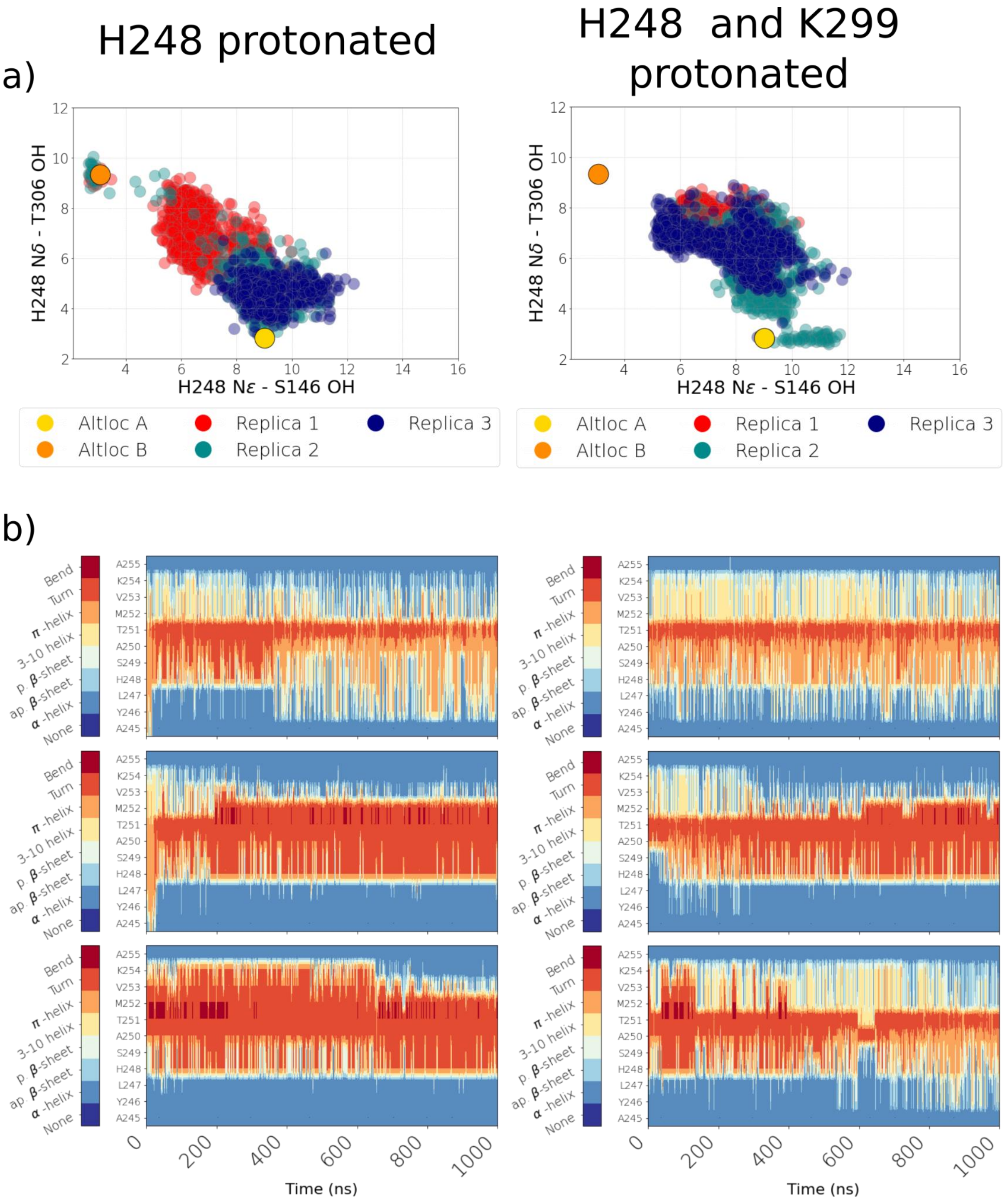

**Figure S24. Dynamics of histidine sidechain and backbone carrying it.** Data from wild type simulation setup number 45. **(a)** Scatter plots display distances of S146 and T306 side chains from H248 side chain. Data is shown from all three simulation replicas in three different colors. Left – H248δ and Right – H248ε **(b)** Time resolved secondary structure analysis of the H248 carrying segment A245-A255. The three rows represent three different simulation replicas. Left – H248δ and Right – H248ε. See main text Fig. 3 for more details.

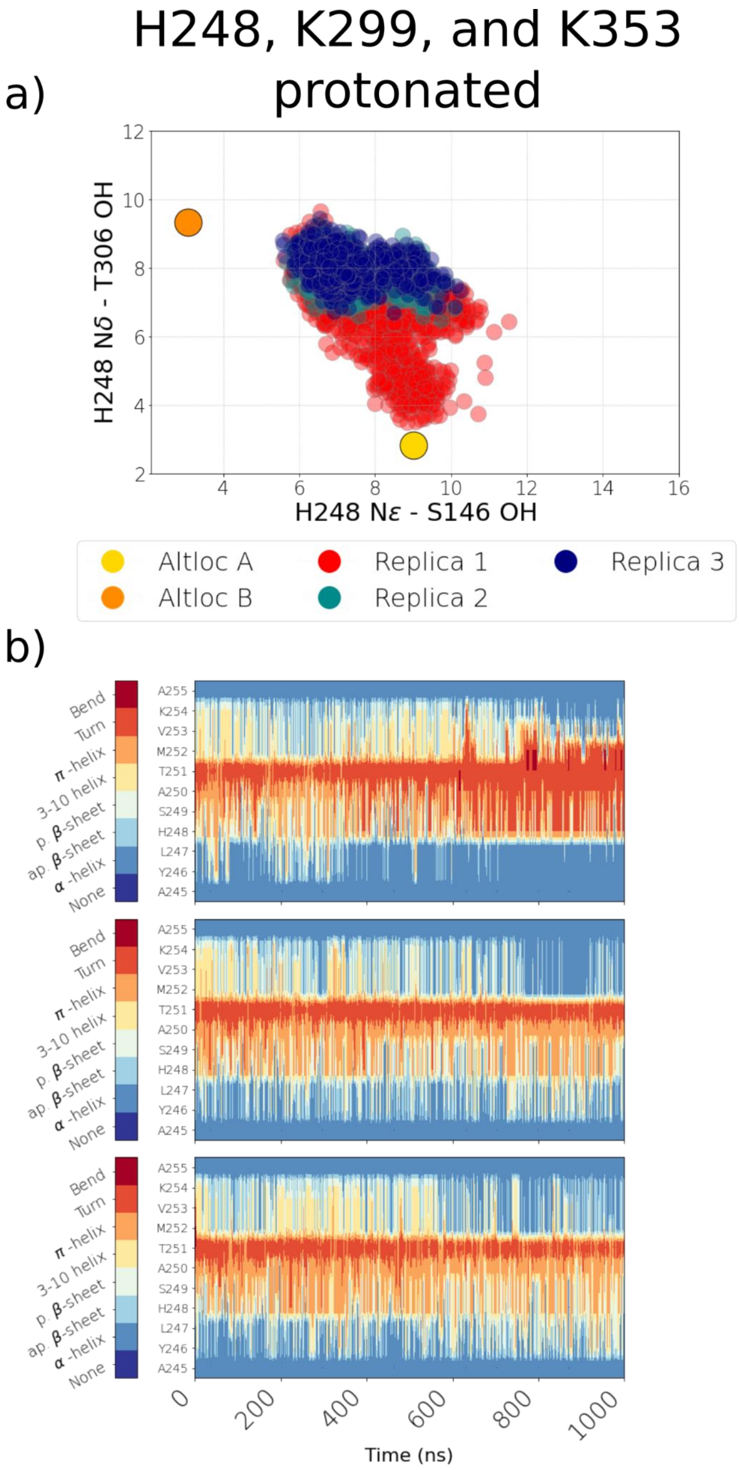

**Figure S25. Dynamics of histidine sidechain and backbone carrying it.** Data from wild type simulation setups number 46 and 47. **(a)** Scatter plots display distances of S146 and T306 side chains from H248 side chain. Data is shown from all three simulation replicas in three different colors. Left – H248δ and Right – H248ε **(b)** Time resolved secondary structure analysis of the H248 carrying segment A245-A255. The three rows represent three different simulation replicas. Left – H248δ and Right – H248ε. See main text Fig. 3 for more details.

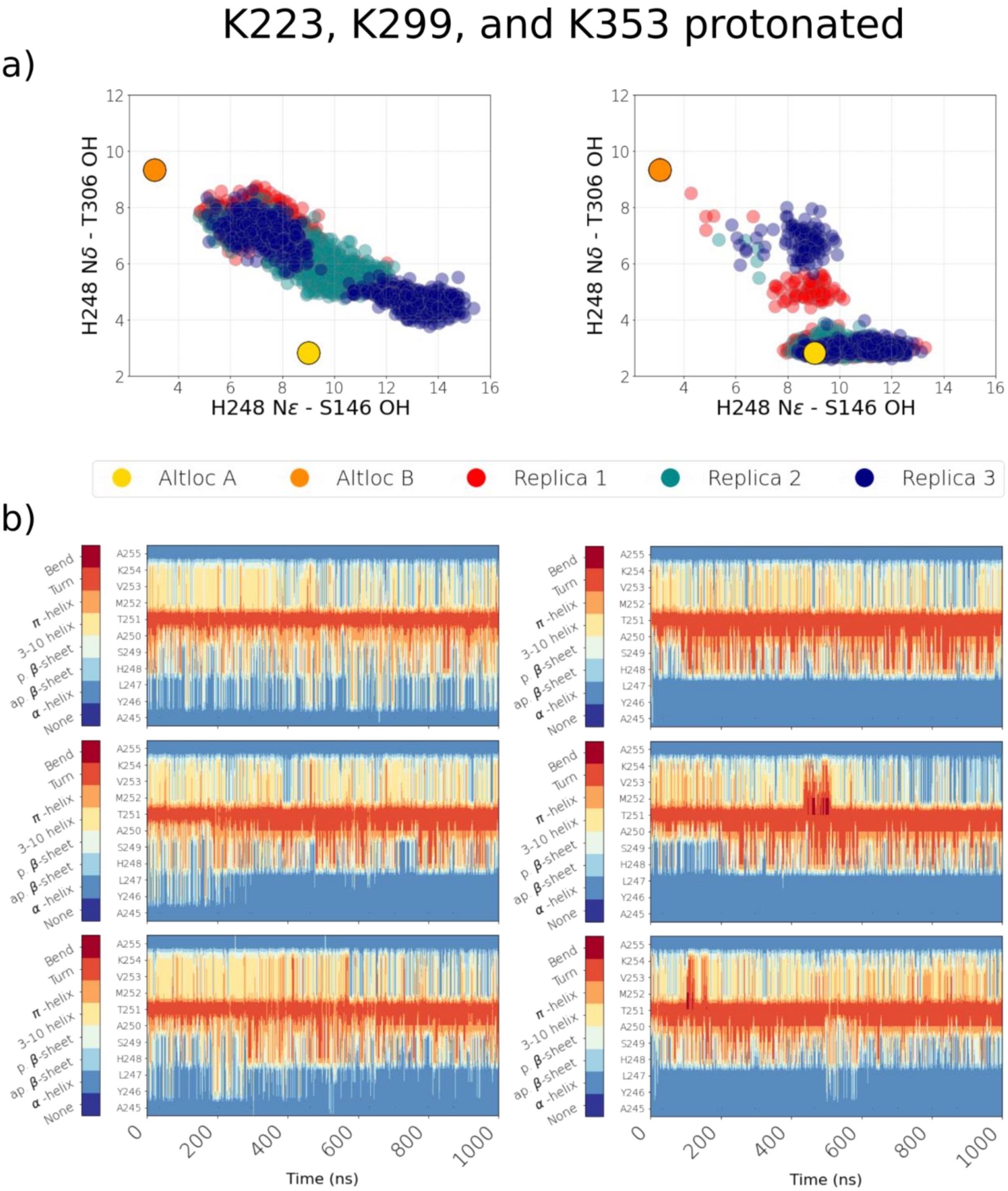

**Figure S26. W232 and alanine mutation, intermediate state from AlphaFold, and change in hydration after mutation of H248 to Ala and Phe. (a)** Comparison between wild type in alternative location A (yellow), and W232A mutation (cyan, representative snapshot). **(b)** Comparison between cryo-EM alternative location A (yellow), alternative location B (orange), H248 protonated (dark red, representative snapshot), and Alpha Fold prediction for MrpA from *Bacillus pseudofirmus* (teal). **(c)** Effect of H248A and **(d)** H248F mutations on hydration. Water occupancy is calculated over a total of 3 µs trajectory data using oxygen atoms of water molecules and is shown at an isovalue of 0.2 (20% occupancy).

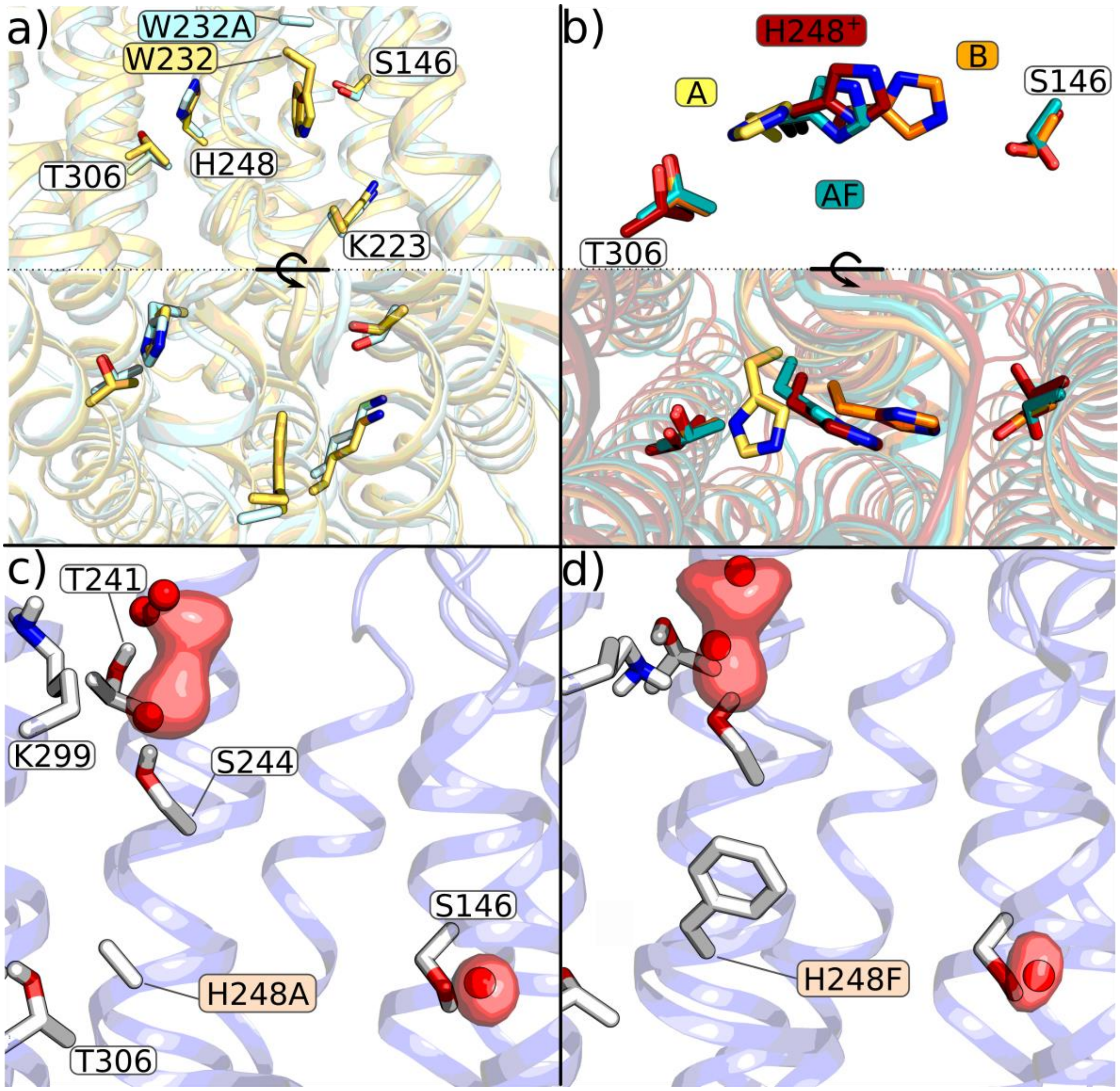

**Figure S27. Loss of helicity in H248 carrying transmembrane helix.** Simulation snapshots displaying backbone hydrogen bonding scenario in a clean alpha helix **(a)** and in the sequence stretch around H248 where alpha helicity is partly lost **(b** and **c)**. In panel b) we show a representative snapshot of A conformation secondary structure rearrangement. Panel c) shows a representative snapshot of B conformation secondary structure rearrangement.

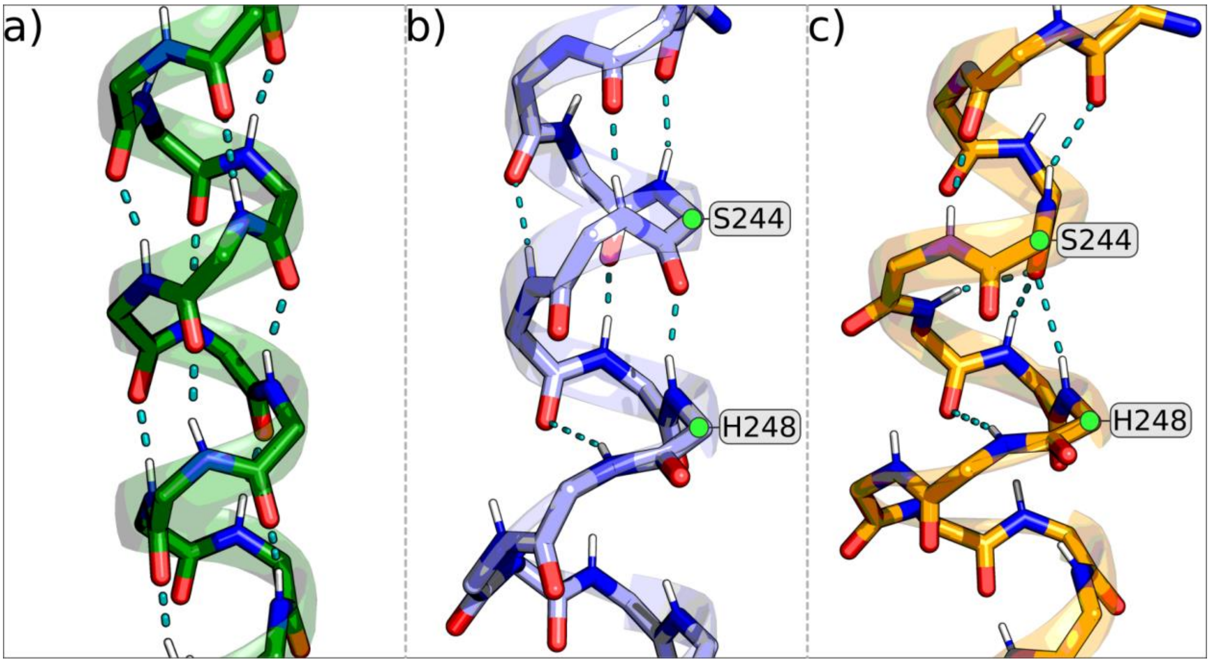

**Figure S28. Protonation-state dependent hydration and conformational changes. a)** H248ε stabilized in B conformation by hydrogen bonding to S146 and a stable water molecule. **b)** Protonated H248 assumes intermediate position and enhances hydration in the region leading to water-based connections to K299 and K223. **c)** Water-based connectivity breaks when K299 is protonated. Water occupancy map shown in the red glass isosurface is calculated over a total of 3 µs trajectory data using oxygen atoms of water molecules and is shown at an isovalue of 0.2 (20% occupancy).

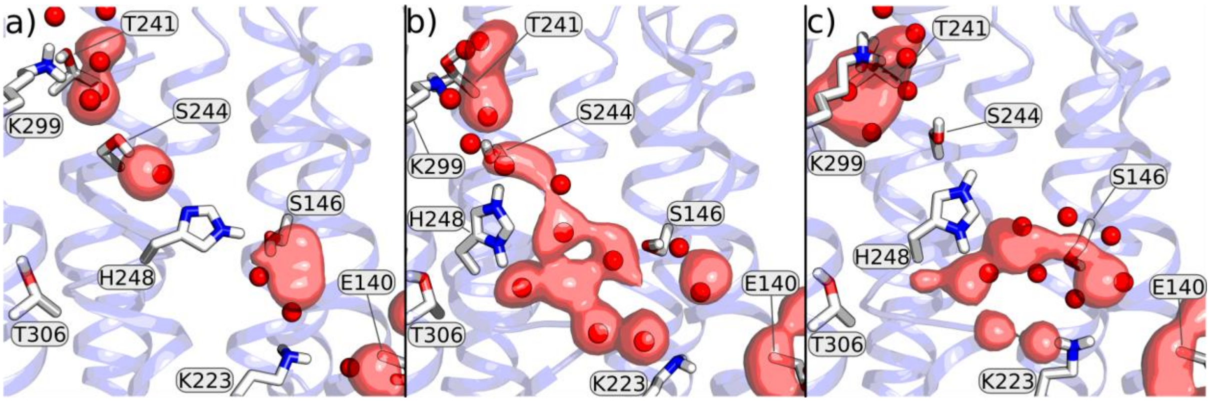

**Figure S29. Proposed histidine switch mechanism of proton transport in ND5 subunit of respiratory complex I.**

All amino acids discussed in the mechanistic proposal shown in Fig. 6 are also conserved in the complex I ND5 subunit. We start with a scenario in which ND5 is initially proton deficient due to the preceding pump cycle. This provides the driving force for proton uptake from the N phase of the membrane by having p$K_a$ of a residue or a group of residues greater than the pH of the N medium. The pre-existing proton hole on K353 and protonated K299 drives the neutral H248 to attain its intermediate position resulting in its protonation from the latter residue (state 1 to 2). Next, K299 gets re-protonated from the cytoplasm or mitochondrial matrix, causing protonated H248 to bind in the A conformation, in part due to the electrostatic repulsion from protonated K299 (state 3). After proton transfer to K353, H248 changes its position to B conformation upon arrival of proton on the Lys/Glu pair of ND5 subunit from the neighboring antiporter-like ND4 subunit (state 4 to 5) (see also simulations of related states in ref. 22). Upon protonation of H248 from the Lys/Glu pair (protonation also possible when H248 is in intermediate position, see main text), the doubly protonated histidine re-assumes the intermediate position (state 6). This high energy state relaxes by pumping of a proton to the P side of the membrane (state 6 to 7) via the ND5 exit (ref. 24), followed by intra-subunit protonic rearrangements. Overall, proton-deficiency in the ND5 subunit drives the uptake of proton from the N phase of the membrane, whereas pumping of the proton to the P side is triggered by the electrostatic push from proton injected by Q redox chemistry (refs. 22 and 36).

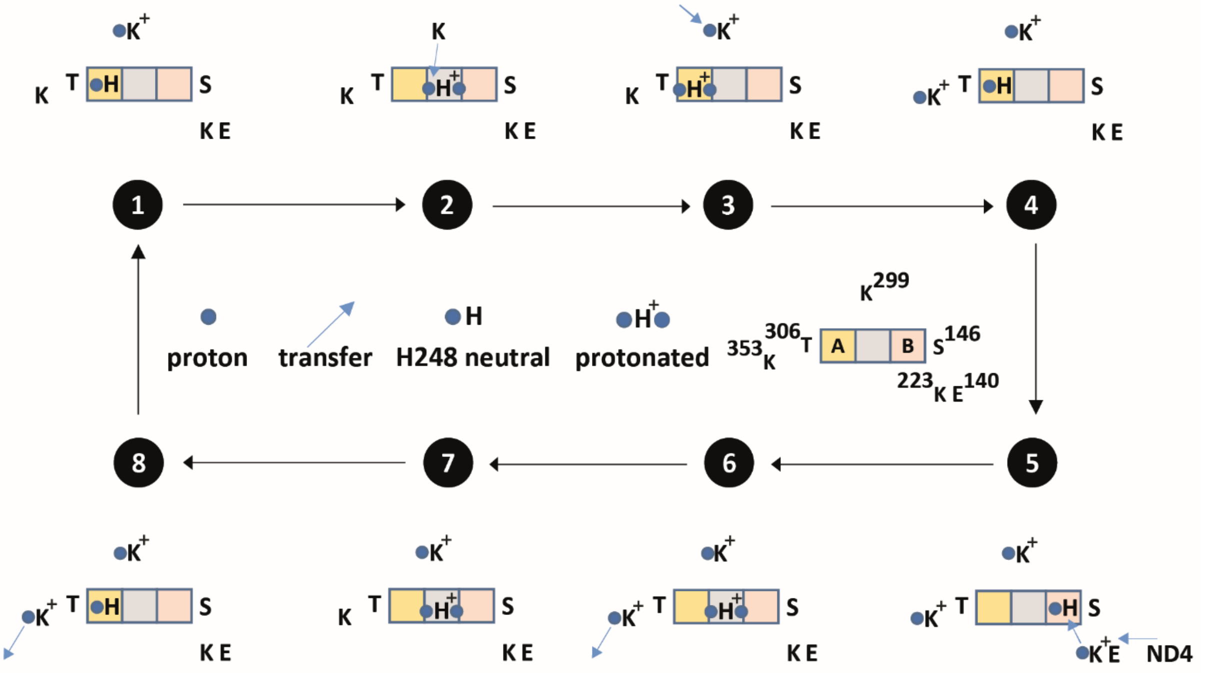

**Figure S30.** A and B conformational occupancies calculated for wild type and mutant conditions. A conformation is adopted when the distance between H248-Nδ and T306-Cα is below 3.75 Å, whereas B conformation is adopted when the distance between H248-Nε and S146-Cα is below 4.75 Å. Percentages are calculated over data from all three simulation replicas of each setup.

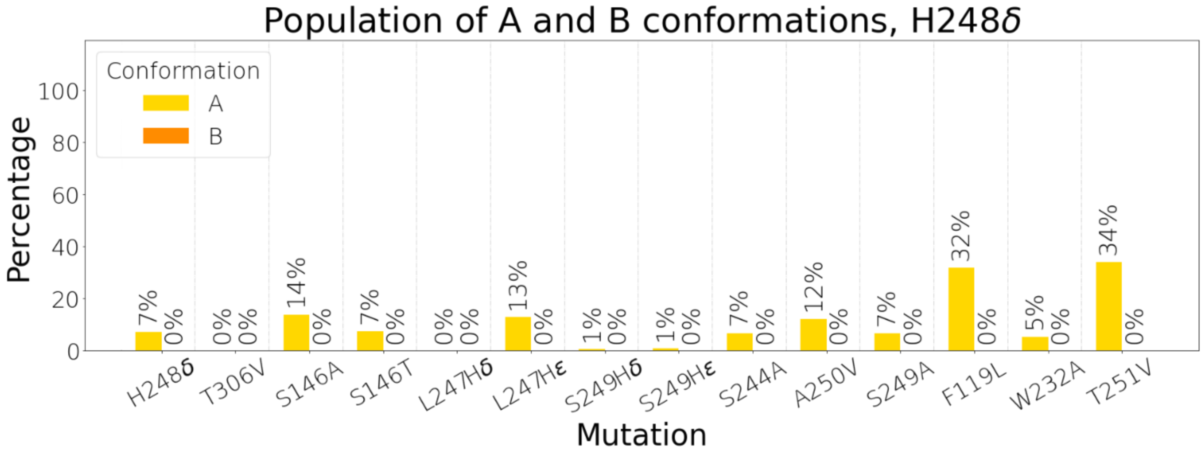

**Figure S31.** A and B conformational occupancies calculated in different charged states of K223, K299, and K353. In H248δ state, all three lysine residues are charge neutral. A conformation is adopted when the distance between H248-Nδ and T306-Cα is below 3.75 Å, whereas B conformation is adopted when the distance between H248-Nε and S146-Cα is below 4.75 Å. Percentages are calculated over data from all three simulation replicas of each setup.

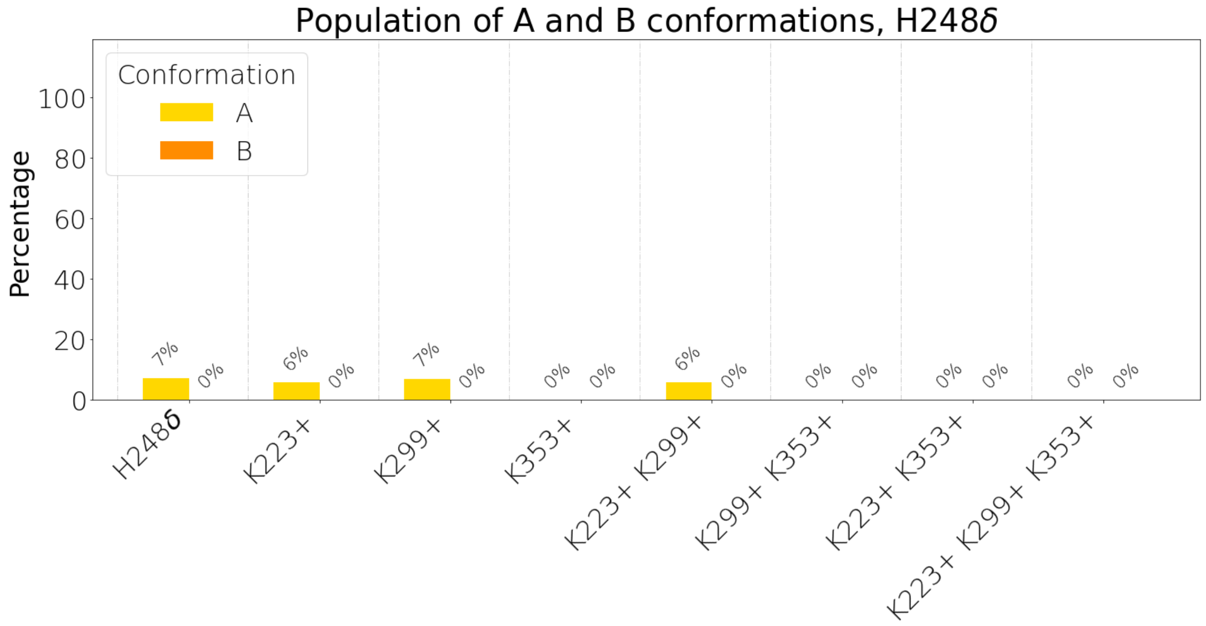

**Figure S32. Snapshots showing comparison between L247H and S249H mutants. (a)** L247Hδ mutant and H248ε. Snapshot displays H248 is occupying A conformation, while L247H interacts with S146. Upper panel is view from the side (see Figure 1), whereas lower panel is view from the cytoplasmic side and provides a zoomed in view. **(b)** S249Hδ mutant and H248ε. Snapshot representing how H248 is occupying B conformation, while S249 points towards nearly the opposite direction. Upper panel is a side view, and lower panel is a view from the cytoplasmic side (zoomed in). Water occupancy map shown in the red glass isosurface is calculated over a total of 3 µs trajectory data using oxygen atoms of water molecules and is shown at an isovalue of 0.2 (20% occupancy). In lower panels, the blue dotted lines highlight a water density near to S249H, which is not seen in L247H simulations.

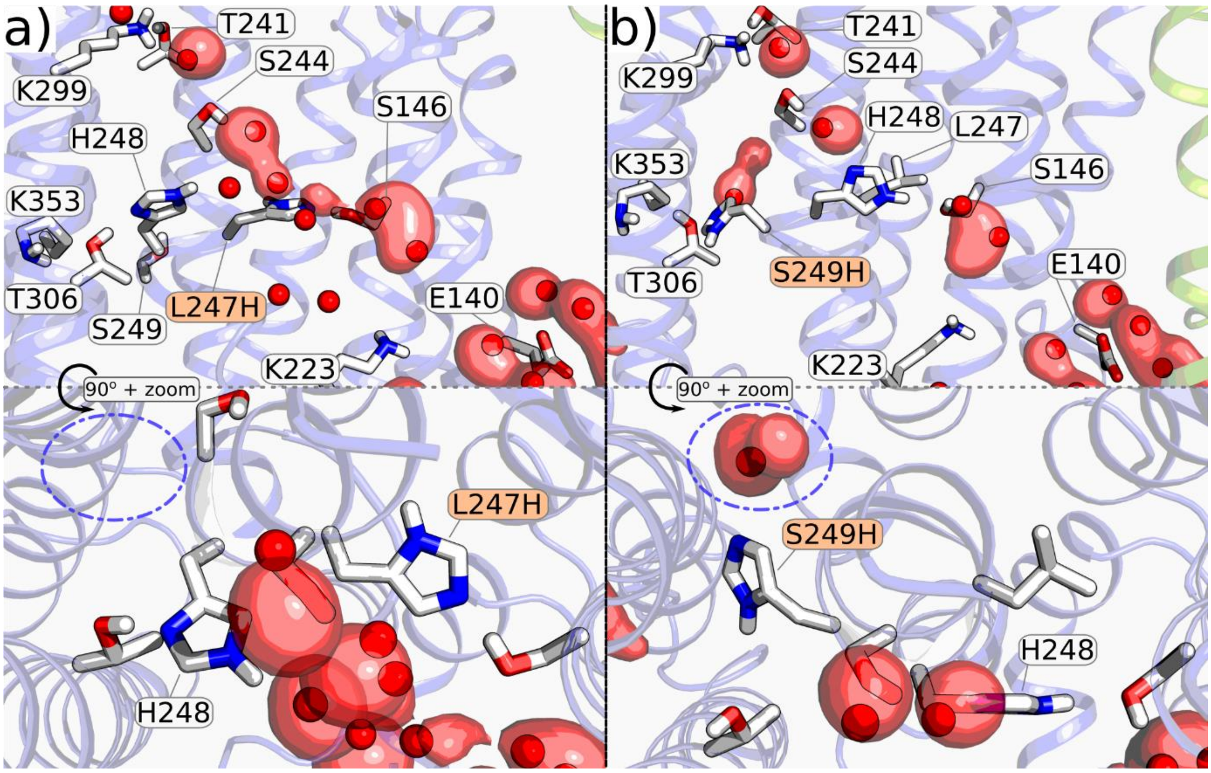

**Figure S33.** Data from simulation setups number 48 and 49. **(a)** Scatter plots display distances of S146 and T306 side chains from H248 side chain. Data is shown from all three simulation replicas in three different colors (simulations are 600 ns). Left – mutation S146A with H248ε and protonated K299 and Right – mutation T306V with H248ε and protonated K353. **(b)** A and B conformational occupancies calculated for the two mutants and corresponding wild type. A conformation is adopted when the distance between H248-Nδ and T306-Cα is below 3.75 Å, whereas B conformation is adopted when the distance between H248-Nε and S146-Cα is below 4.75 Å. Percentages are calculated over data from all three simulation replicas of each setup. Because these mutant MD simulations are 600 ns long, here we show data from the first 600 ns of wild type MD simulations.

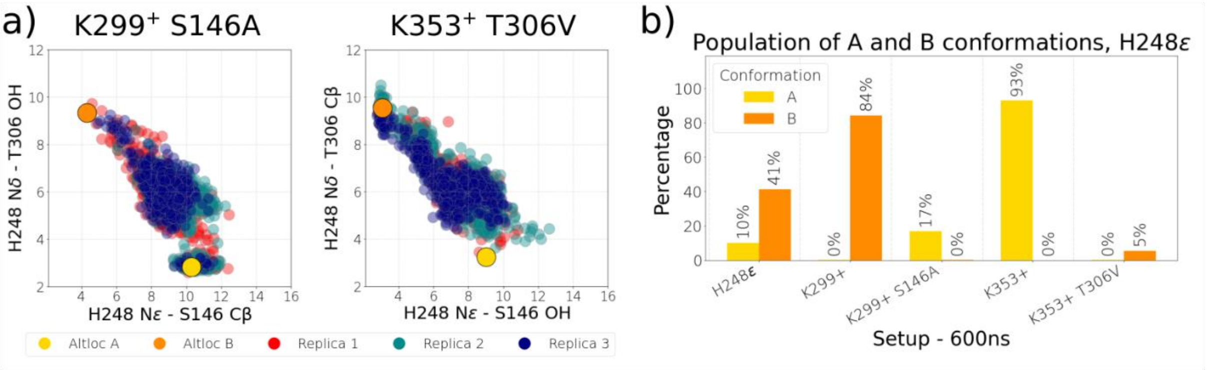

**Table S1. Strains and Plasmids used in this study.**

| Strain - plasmid | Description - Genotype | Source |
|---|---|---|
| *Escherichia coli* Strains | | |
| KNabc | TG1(Δ*nhaA* Δ*nhaB* Δ*chaA*) | (29) |
| KNabc(DE3) | KNabc (DE3) | This Work |
| XL10-Gold | K12 ($Tet^r$ Δ(*mcrA*)*183* Δ(*mcr*CB-*hsdSMR*-*mrr*)*173* *endA1* *supE44* *thi-1* *recA1* *gyrA96*($nal^R$) *relA1* *lac* Hte) F' [*proAB* *lacI^q*ZΔ*M15* Tn*10*($Tet^r$ Amy $Cam^r$)] | Stratagene |
| Plasmids | | |
| pUB26 | Empty cloning plasmid conferring Ampelicillin-resistance | Kerscher et al. 2002 |
| pGEM_BpMrp_WT | pGEM expression plasmid conferring Ampelicillin-resistance and carrying the full Mrp operon from *Bacillus pseudofirmus* with a native up stream sequence of 170 bp. The Gen for MrpG carries a $His_6$-FLAG tag | (19) |
| pGEM_BpMrp_V2_WT | pGEM_BpMrp_WT plasmid without the 170 bp up stream sequence and silent mutations in MrpA at Val57, Val58 and Ala122 | This Work |
| pGEM_BpMrp_V2_MrpA-F119L | pGEM_BpMrp_V2_WT + mutant MrpA-F119L | This Work |
| pGEM_BpMrp_V2_MrpA-S146A | pGEM_BpMrp_V2_WT + mutant MrpA-S146A | This Work |
| pGEM_BpMrp_V2_MrpA-S146T | pGEM_BpMrp_V2_WT + mutant MrpA-S146T | This Work |
| pGEM_BpMrp_V2_MrpA-S146V | pGEM_BpMrp_V2_WT + mutant MrpA-S146V | This Work |
| pGEM_BpMrp_V2_MrpA-W232A | pGEM_BpMrp_V2_WT + mutant MrpA-W232A | This Work |
| pGEM_BpMrp_V2_MrpA-W232Y | pGEM_BpMrp_V2_WT + mutant MrpA-W232Y | This Work |
| pGEM_BpMrp_V2_MrpA-S244A | pGEM_BpMrp_V2_WT + mutant MrpA-S244A | This Work |
| pGEM_BpMrp_V2_MrpA-S244V | pGEM_BpMrp_V2_WT + mutant MrpA-S244V | This Work |

| | | |
|---|---|---|
| pGEM_BpMrp_V2_MrpA-L247H | pGEM_BpMrp_V2_WT + mutant MrpA-L247H | This Work |
| pGEM_BpMrp_V2_MrpA-H248A | pGEM_BpMrp_V2_WT + mutant MrpA-H248A | This Work |
| pGEM_BpMrp_V2_MrpA-H248F | pGEM_BpMrp_V2_WT + mutant MrpA-H248F | This Work |
| pGEM_BpMrp_V2_MrpA-S249A | pGEM_BpMrp_V2_WT + mutant MrpA-S249A | This Work |
| pGEM_BpMrp_V2_MrpA-S249H | pGEM_BpMrp_V2_WT + mutant MrpA-S249H | This Work |
| pGEM_BpMrp_V2_MrpA-A250C | pGEM_BpMrp_V2_WT + mutant MrpA-A250C | This Work |
| pGEM_BpMrp_V2_MrpA-A250S | pGEM_BpMrp_V2_WT + mutant MrpA-A250S | This Work |
| pGEM_BpMrp_V2_MrpA-A250V | pGEM_BpMrp_V2_WT + mutant MrpA-A250V | This Work |
| pGEM_BpMrp_V2_MrpA-T251V | pGEM_BpMrp_V2_WT + mutant MrpA-T251V | This Work |
| pGEM_BpMrp_V2_MrpA-T306A | pGEM_BpMrp_V2_WT + mutant MrpA-T306A | This Work |
| pGEM_BpMrp_V2_MrpA-T306S | pGEM_BpMrp_V2_WT + mutant MrpA-T306S | This Work |
| pGEM_BpMrp_V2_MrpA-T306V | pGEM_BpMrp_V2_WT + mutant MrpA-T306V | This Work |

**Table S2. Primers used in this study.**

| Primer | Sequence (5’-3’) |
|---|---|
| ΔnatPromotor_for | CATATGagtACCGTATTACATTGGGCTAC |
| ΔnatPromotor_rev | ACGGTactCATATGTATATCTCCCGAATTCCC |
| ΔTN10#1_for | GGGCgATGtTTGGAGTTGTCTTATCAGATAAC |
| ΔTN10#1_rev | TCCAAaCATcGCCCCCATGAACATGAG |
| ΔTN10#2_for | GGGTgGTgGAGCATACGATTCCATGGG |
| ΔTN10#2_rev | TATGCTCcACcACCCCTCCAGTTGAGG |
| MrpA-F119A_for | CTCATGgcgATGGGGGCGATGTTTG |
| MrpA-F119A_rev | CCCCCATcgcCATGAGAAGGTACACATAGAAAT TG |
| MrpA-S146A_for | GCAgcgTCGCTATTAATTAGTTATTGGTTCC |
| MrpA-S146A_rev | GCGAcgcTGCTAAACTAGTTAATTCCCAAAAC |
| MrpA-S146T_for | AGCAaccTCGCTATTAATTAGTTATTGGTTCC |
| MrpA-S146T_rev | GCGAggtTGCTAAACTAGTTAATTCCCAAAAC |
| MrpA-S146V_for | AGCAgtgTCGCTATTAATTAGTTATTGGTTCC |
| MrpA-S146V_rev | GCGAcacTGCTAAACTAGTTAATTCCCAAAAC |
| MrpA-W232A_for | ACATCgcGCTGCCAGATGCCATG |
| MrpA-W232A_rev | CAGCgcGATGTGGAAGGGGAACTG |
| MrpA-W232Y_for | ACATCTatCTGCCAGATGCCATG |
| MrpA-W232Y_rev | GGCAGatAGATGTGGAAGGGGAAC |
| MrpA-S244A_for | ACCTGTTgcgGCATATTTACACTCTGCAAC |
| MrpA-S244A_rev | AAATATGCcgcAACAGGTGTTGGTGCTTC |
| MrpA-S244V_for | ACCTGTTgtgGCATATTTACACTCTGCAAC |
| MrpA-S244V_rev | AAATATGCcacAACAGGTGTTGGTGCTTC |
| MrpA-L247H_for | GCATATcatCACTCTGCAACAATGGTTAAAG |
| MrpA-L247H_rev | AGAGTGatgATATGCACTAACAGGTGTTGG |
| MrpA-H248A_for | GCATATTTAgcgTCTGCAACAATGGTTAAAGC |
| MrpA-H248A_rev | GCAGAcgcTAAATATGCACTAACAGGTGTTG |
| MrpA-H248F_for | GCATATTTAtttTCTGCAACAATGGTTAAAGC |
| MrpA-H248F_rev | GCAGAaaaTAAATATGCACTAACAGGTGTTG |
| MrpA-S249A_for | ACACgcgGCAACAATGGTTAAAGCAG |
| MrpA-S249A_rev | GTTGCcgcGTGTAAATATGCACTAACAGG |
| MrpA-S249H_for | TTACACcatGCAACAATGGTTAAAGCAG |
| MrpA-S249H_rev | TGCatgGTGTAAATATGCACTAACAGGTG |
| MrpA-A250C_for | CACTCTtgcACAATGGTTAAAGCAGGGATCT |
| MrpA-A250C_rev | CCATTGTgcaAGAGTGTAAATATGCACTAACAG G |
| MrpA-A250S_for | CACTCTagcACAATGGTTAAAGCAGGGATCT |
| MrpA-A250S_rev | CCATTGTgctAGAGTGTAAATATGCACTAACAGG |
| MrpA-A250V_for | CTCTGtgACAATGGTTAAAGCAGGGATCT |
| MrpA-A250V_rev | CATTGTcaCAGAGTGTAAATATGCACTAACAGG |

| | |
|---|---|
| MrpA-T251V_for | TCTGCAgtgATGGTTAAAGCAGGGATCTATC |
| MrpA-T251V_rev | AACCATcacTGCAGAGTGTAAATATGCACTAAC |
| MrpA-T306A_for | CTCAgcgGTCAGTCAGCTCGGTCTG |
| MrpA-T306A_rev | ACTGACcgcTGAGAAGGCGAGAATTCC |
| MrpA-T306S_for | CTCAAgcGTCAGTCAGCTCGGTCTG |
| MrpA-T306S_rev | TGACgcTTGAGAAGGCGAGAATTCC |
| MrpA-T306V_for | CTCAgtgGTCAGTCAGCTCGGTCTG |
| MrpA-T306V_rev | ACTGACcacTGAGAAGGCGAGAATTCC |

**Table S3. Protonation states based on Propka analysis (see computational methods)**

| Subunit | Residue | Charge state A conformation (pKa) | Charge state B conformation (pKa) |
|---|---|---|---|
| MrpA | Lys223 | 0 (6.94) | 0 (6.65) |
| MrpA | Lys254 | 0 (5.85) | 0 (5.90) |
| MrpA | Lys299 | 0 (6.86) | 0 (6.84) |
| MrpA | Lys353 | 0 (6.40) | 0 (6.43) |
| MrpA | Lys408 | +1 (7.94) | 0 (6.37) |
| MrpA | Glu409 | 0 (7.20) | -1 (6.79) |
| MrpA | His470 | +1 (7.17) | +1 (7.17) |
| MrpA | Asp678 | 0 (7.24) | 0 (7.36) |
| MrpA | Glu687 | 0 (8.42) | 0 (7.75) |
| MrpA | Asp771 | 0 (7.65) | 0 (7.65) |
| MrpA | Glu780 | 0 (8.54) | 0 (9.52) |
| MrpB | Asp121 | 0 (7.34) | 0 (7.35) |
| MrpD | Lys250 | 0 (6.31) | 0 (6.31) |
| MrpD | Lys337 | 0 (5.61) | 0 (5.61) |
| MrpF | Asp38 | -1 (6.70) | 0 (7.03) |

**Table S4.** MD simulation setups. Each setup was simulated in triplicate, with each simulation replica 1000 ns long unless otherwise mentioned. Note, all H248δ state simulations were initiated from A alternative conformation, whereas H248ε simulations from the B alternative conformation. * - Simulations length 500 ns. ** - Simulations length 600 ns.

| Setup number | Setup | Protonation state of selected residues (all other titratable residues are in the charge state determined by pKa calculations, see Table S3) | Starting conformation altloc |
|---|---|---|---|
| 1 | Wild type | H248δ | A |
| 2 | Wild type | H248ε | B |
| 3 | Wild type | $H248^{+}$ | B |
| 4 | F119L | H248δ | A |
| 5 | F119L | H248ε | B |
| 6 | S146A | H248δ | A |
| 7 | S146A | H248ε | B |
| 8 | S146T | H248δ | A |
| 9 | S146T | H248ε | B |
| 10 | W232A | H248δ | A |
| 11 | W232A | H248ε | B |
| 12 | S244A | H248δ | A |
| 13 | S244A | H248ε | B |
| 14 | L247Hδ | H248δ | A |
| 15 | L247Hδ | H248ε | B |
| 16 | L247Hε | H248δ | A |
| 17 | L247Hε | H248ε | B |
| 18 | H248A* | - | A |
| 19 | H248F* | - | A |
| 20 | S249A | H248δ | A |
| 21 | S249A | H248ε | B |
| 22 | S249Hδ | H248δ | A |
| 23 | S249Hδ | H248ε | B |
| 24 | S249Hε | H248δ | A |
| 25 | S249Hε | H248ε | B |

| | | | |
|---|---|---|---|
| 26 | A250V | H248δ | A |
| 27 | A250V | H248ε | B |
| 28 | T251V | H248δ | A |
| 29 | T251V | H248ε | B |
| 30 | T306V | H248δ | A |
| 31 | T306V | H248ε | B |
| 32 | Wild type | H248δ K223+ | A |
| 33 | Wild type | H248ε K223+ | B |
| 34 | Wild type | H248δ K299+ | A |
| 35 | Wild type | H248ε K299+ | B |
| 36 | Wild type | H248δ K353+ | A |
| 37 | Wild type | H248ε K353+ | B |
| 38 | Wild type | H248δ K223+ K299+ | A |
| 39 | Wild type | H248ε K223+ K299+ | B |
| 40 | Wild type | H248δ K223+ K353+ | A |
| 41 | Wild type | H248ε K223+ K353+ | B |
| 42 | Wild type | H248+ K299+ | A |
| 43 | Wild type | H248δ K299+ K353+ | A |
| 44 | Wild type | H248ε K299+ K353+ | B |
| 45 | Wild type | H248+ K299+ K353+ | A |
| 46 | Wild type | H248δ K223+ K299+ K353+ | A |
| 47 | Wild type | H248ε K223+ K299+ K353+ | B |
| 48 | T306V** | H248ε K353+ | B |
| 49 | S146A** | H248ε K299+ | B |
